# Paths to Oblivion: Common Neural Mechanisms of Anaesthesia and Disorders of Consciousness

**DOI:** 10.1101/2021.02.14.431140

**Authors:** Andrea I. Luppi, Pedro A.M. Mediano, Fernando E. Rosas, Judith Allanson, John D. Pickard, Guy B. Williams, Michael M Craig, Paola Finoia, Alexander R.D. Peattie, Peter Coppola, Adrian Owen, Lorina Naci, David K. Menon, Daniel Bor, Emmanuel A. Stamatakis

## Abstract

The human brain generates a rich repertoire of spatiotemporal dynamics during normal wakefulness, supporting a wide variety of conscious experiences and cognitive functions. However, neural dynamics are reconfigured, in comparable ways, when consciousness is lost either due to anaesthesia or disorders of consciousness (DOC). Here, leveraging a neurobiologically realistic whole-brain computational model informed by functional MRI, diffusion MRI, and PET, we sought to identify the neurobiological mechanisms that explain the common reconfiguration of neural dynamics observed both for transient pharmacological intervention and chronic neuroanatomical injury. Our results show that, by incorporating local inhibitory action through a PET-based GABA receptor density map, our model can reproduce the brain dynamics of subjects undergoing propofol anaesthesia, and that this effect depends specifically on the spatial distribution of GABA receptors across cortical regions. Additionally, using a structural connectome obtained from DOC patients, we demonstrate how the dynamics that characterise loss of consciousness can emerge from changes in neuroanatomical connectivity. Crucially, we find that each of these two interventions generalises across datasets: a model with increased GABA-mediated inhibition can reproduce the dynamics of DOC patients’ brains, and a model with a DOC connectome is also compatible with brain dynamics observed during propofol anaesthesia. These results demonstrate how increased inhibition and connectome randomisation represent different neurobiological paths towards the characteristic dynamics of the unconscious brain. Overall, the present findings begin to disentangle the neurobiological mechanisms by which highly dissimilar perturbations of the brain’s neurodynamics can lead to unconsciousness.

## Introduction

A central challenge of contemporary neuroscience is the quest to understand how the neurobiology and function of the human brain give rise to conscious experience ^1,2^. One way to address this question is to identify changes in brain function that accompany changes in conscious state. However, the brain is a paradigmatic example of a complex system ^3^, and different perturbations of its precise functioning can serve as a path towards loss of consciousness. Examples of such perturbations range from transient pharmacological (general anaesthetic) interventions having widespread effects on neuromodulation ^4–7^, to chronic disorders of consciousness arising from traumatic or anoxic injuries of diverse location and extent, often including changes to the physical connectivity between brain regions ^8–12^.

In order to shed light on how the human brain supports consciousness, two fundamental challenges must be addressed. The first is to identify neurobiological signatures of consciousness and its loss that are generalisable across different paths to unconsciousness, rather than being specific to any of them ^13^. The second is to provide a mechanistic understanding of the previously identified signatures of consciousness, in terms of the underlying neurobiological principles ^14^.

Substantial progress has recently been made towards answering the first question. The human brain generates a constantly changing repertoire of neural dynamics, supporting the rich variety of conscious experiences and cognitive functions ^15,16,25–29,17–24^. These complex dynamics self-organise ^30^ according to the recursive local-global interplay of neuronal excitation and inhibition, and their intricate interactions across the network of the brain’s anatomical connections (human connectome) ^16,31–37^.

Recently, converging evidence from multimodal neuroimaging has further shown that brain dynamics are substantially and consistently altered when consciousness is lost, which suggests that specific aspects of neural dynamics may play a central role in supporting human consciousness. In particular, striking similarities have been observed across brain dynamics induced by different ways of losing consciousness, such as acute pharmacological intervention with different anaesthetic drugs, but also chronic disorders of consciousness ^13,38–44^. These findings suggest that spatio-temporal dynamics may provide a “common currency” between brain and mind ^45^ capable of encoding the difference between consciousness and unconsciousness.

Despite recent achievements in finding common signatures of consciousness and its loss, the causal neurobiological mechanisms underlying these processes remain elusive. For example, anaesthesia is known to operate at the level of neurotransmission (propofol is a potent agonist of inhibitory GABA-A receptors ^46,47^), without affecting the physical organisation of the human connectome. In contrast, disorders of consciousness (DOC) arise from severe brain injury, typically due to anoxia or head trauma, which cause brain lesions of variable severity and location often including alterations of the physical connectivity between patients’ brain regions ^10,48–56^. This similarity of outcomes and neural signatures ^13,38–42^ despite arising from radically different causes, begs the question: How can (transient) pharmacological and (chronic) structural perturbations converge to similar effects on brain dynamics, and the corresponding state of unconsciousness?

Here, we sought to obtain mechanistic insights into this fundamental question by employing whole-brain computational modelling, which is emerging as a powerful tool to investigate the neurobiological mechanisms underlying macroscale neural phenomena ^14,35,57–60^. These models represent regional macroscale activity in terms of local dynamics influenced by inter-regional anatomical connectivity (obtained e.g. from diffusion MRI tractography), using approximations derived from oscillatory mechanisms (e.g. Kuramoto model), statistical mechanics (e.g. Ising model), or dynamical systems theory (e.g. Hopf model) ^35,61^.

Crucially, *in silico* models are uniquely suited to investigate how different neurobiological perturbations can induce similar alterations of brain dynamics ^14^. Just like neuropsychological studies in human patients and experimental lesions in animal models have provided invaluable insights about brain organisation, function and dysfunction ^62–65^, whole-brain computational models can be systematically and reversibly lesioned to investigate the resulting alterations in macroscale brain dynamics. Being fully accessible to the researcher, the model’s parameters can be systematically altered in ways that are still beyond the capabilities of experimental research, whether in humans or animals ^14,66^.

For these reasons, whole-brain computational modelling is becoming increasingly prominent as a tool to investigate the causal mechanisms that drive brain network organisation in healthy and pathological conditions ^35,57,75,67–74^. In particular, recent efforts have capitalised on the tractability of the Hopf and Ising models, beginning to shed light on the mechanisms of loss of consciousness during sleep, anaesthesia, and disorders of consciousness in terms of local-global synchronisation behaviour (Hopf model) ^76–82^ or statistical temperature (Ising model) ^83–85^.

Crucially, recent work has demonstrated that more detailed biophysical models that incorporate neurophysiologically realistic information about excitation, inhibition and neuromodulation – so-called *Dynamic Mean Field* (DMF) models – can provide insights about pharmacologically-induced changes in macroscale fMRI dynamics, in terms of the underlying neurobiology ^86–88^. These models reduce the intricate dynamics of individual neurons to a set of coupled differential equations which approximate the detailed microscale neural properties of spiking neurons (incorporating realistic aspects of neurophysiology such as synaptic dynamics and membrane potential) ^89^ via a mean-field reduction ^31,34,37,90,91^. Specifically, cortical regions are represented as macroscopic neural fields, whose local dynamics are coupled together by a network of anatomical connections ^31,34,37^. An additional biophysical haemodynamic model can then be used to turn the DMF model’s dynamics into a realistic simulator of BOLD signals ^92^. Thus, neurobiologically realistic whole-brain computational modelling provides a principled way to bridge across scales, relating the macroscale neural dynamics of fMRI to the microscale neurophysiological mechanisms from which they emerge ^14^.

However, to date no studies have harnessed the power of DMF models to provide neurobiologically realistic accounts of pharmacological and chronic loss of consciousness. Such an effort is vital because as Sanz-Perl and colleagues remark, “*modeling efforts incorporating more complex dynamics could allow in silico rehearsal of interventions with ampler neurobiological interpretation”* ^80^. Here, we leveraged a neurobiologically realistic DMF model informed by multimodal neuroimaging including empirical dynamics from functional MRI, anatomical connectivity obtained from diffusion MRI, and GABA-A receptor density estimated from positron emission tomography (PET) (Figure 1).

**Figure 1.**
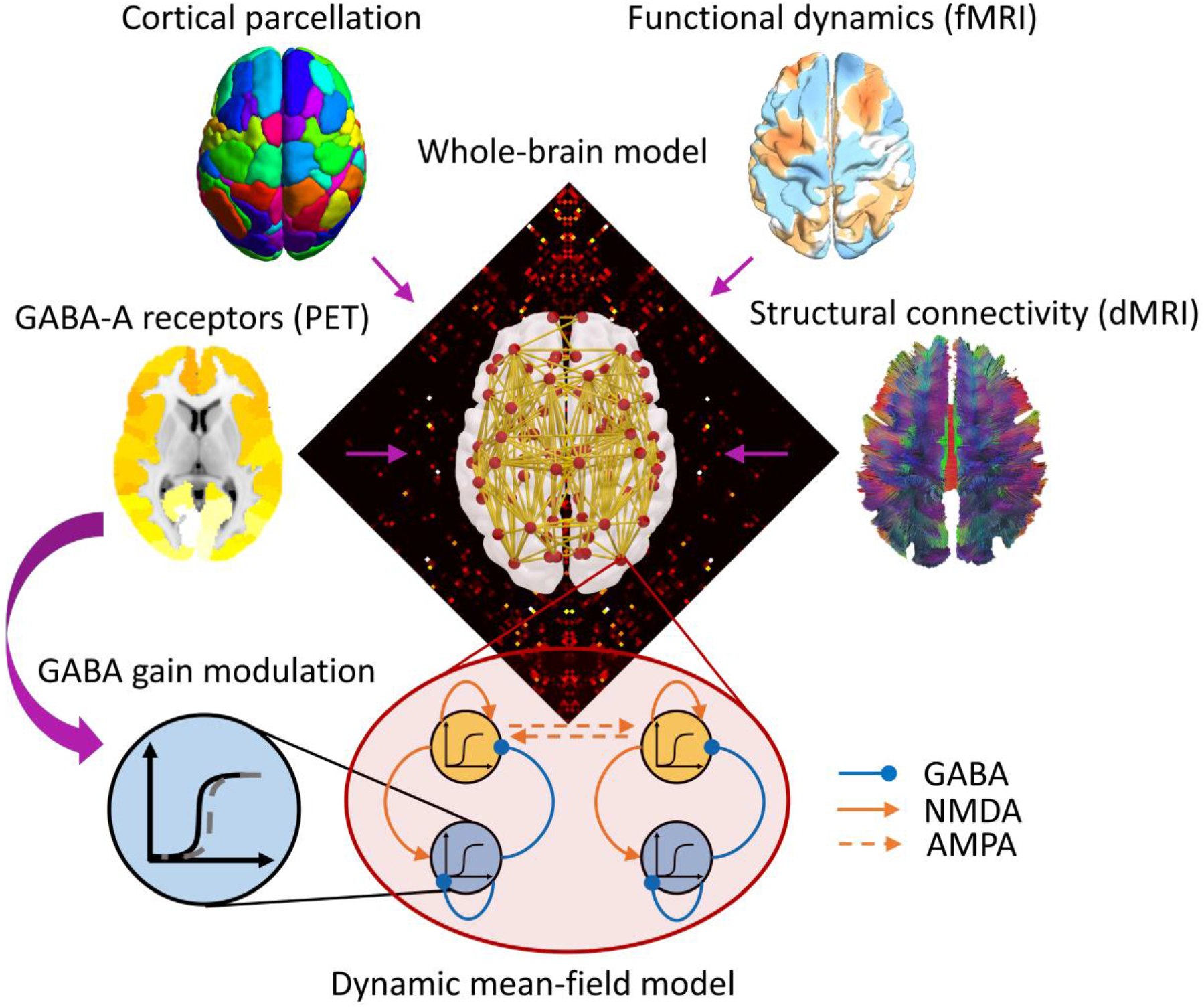
Overview of whole-brain computational model incorporating multimodal neuroimaging data. Based on a cortical parcellation with 68 regions of interest, each node (cortical region) is modelled through a neurophysiologically realistic biophysical model incorporating excitatory (NMDA) as well as inhibitory (GABA) synaptic dynamics. Nodes are connected by structural connectivity (from diffusion MRI) and the model’s dynamics are fitted to simulate empirical brain dynamics (from functional MRI). Neurotransmitter information from PET can also be added in the model as modulating the local neuronal gain ^86^.

We used this modelling approach to simulate the empirical fMRI macroscale brain dynamics observed in the same N=16 subjects at baseline and during loss of consciousness induced by the intravenous anaesthetic, propofol. We also studied the fMRI dynamics of a cohort (N=21) of patients suffering from chronic disorders of consciousness (DOC) as a result of severe brain injury (traumatic or anoxic), comparing them with a group of N=20 healthy controls. By subjecting the models to “virtual anaesthesia” (local modulation of inhibitory gain based on empirical GABA-A receptor distribution) and “virtual DOC” (alteration of the model’s structural connectome), we sought to identify the neurobiological mechanisms underlying a fundamental question of modern neuroscience: how can transient perturbations of neurotransmission and chronic lesions to the structural connectome, both give rise to unconsciousness and its characteristic similar brain dynamics ^13,38,39,41,93^?

## Results

The neurobiologically realistic dynamic mean-field (DMF) model only one free parameter: a global coupling parameter, denoted by *G*, which scales the excitatory-to-excitatory coupling between brain regions, as established by the empirical structural connectome. Thus, calibrating the model corresponds to finding the value of *G* that allows the model to best simulate observed fMRI dynamics of the human brain at rest (which is assessed in terms of the KS-distance between real and simulated data, see ^86^). We followed this procedure for each of our two datasets (Figure 2): for the propofol dataset we optimised the model to fit the fMRI dynamics acquired during the baseline awake scan, and for the DOC dataset we optimised the model to fit the dynamics observed in the healthy controls. These two calibrated DMF models - with their corresponding global coupling values fitted to the dynamics of the conscious brain - constitute the starting point for our investigations.

**Figure 2.**
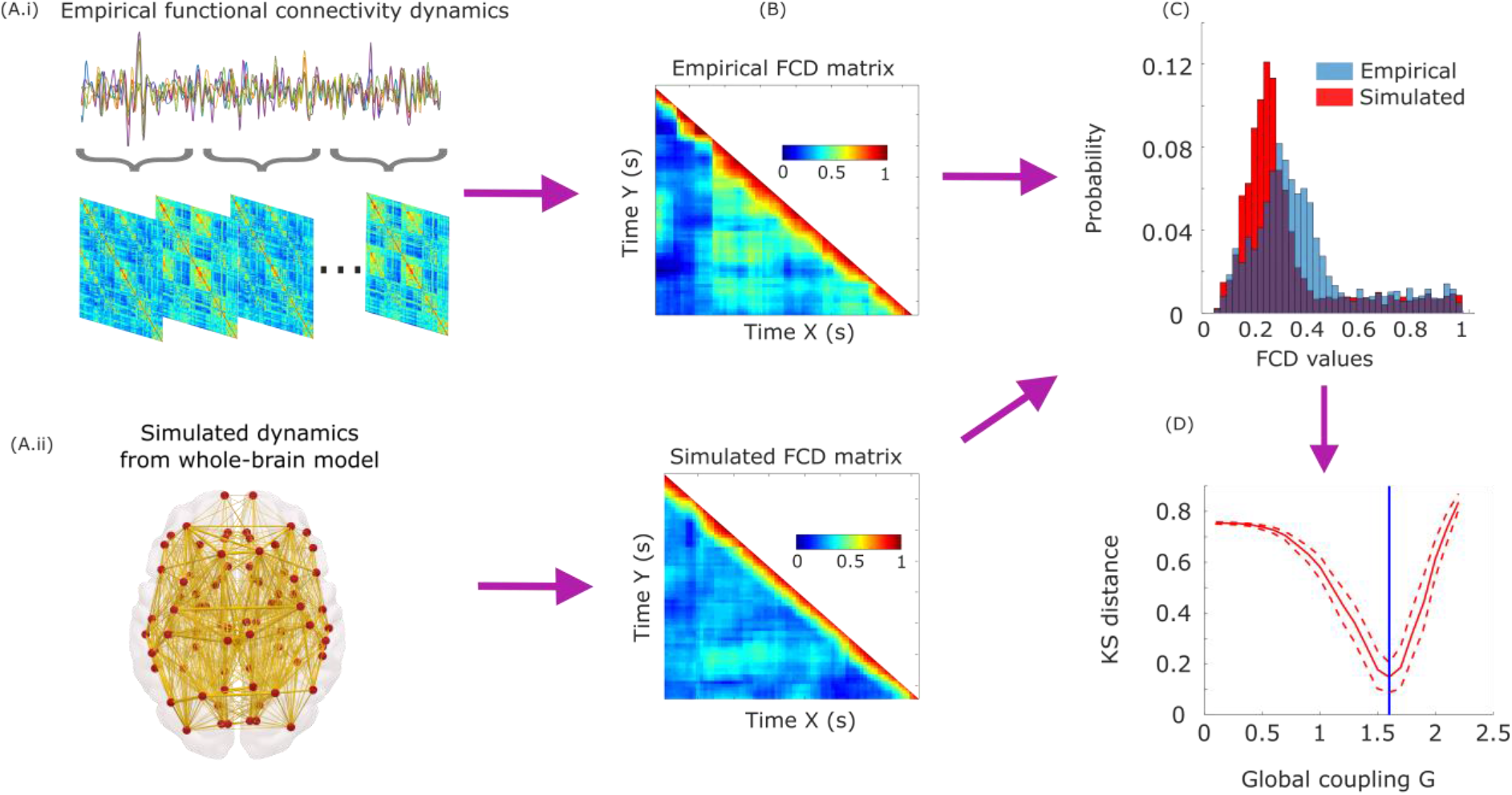
Overview of model fitting procedure. (A.i) Time-resolved matrices of functional connectivity are obtained from empirical functional MRI via the sliding-window approach: regional BOLD time-series are partitioned into windows of 30 TRs, sliding by 3 TRs at a time, following the same approach as previous work using the DMF model ^86^; functional connectivity between each pair of regions is computed within each window by means of Pearson correlation, generating a stack of FC matrices representing the evolution of FC over time. (A.ii) The same procedure is repeated for the simulated BOLD timeseries produced by the model with various levels of the global coupling parameter, *G*. (B) For both the empirical and simulated functional connectivity dynamics (FCD), a time-versus-time FCD matrix is computed by correlating the time-dependent FC matrices centred at each timepoint. (C) Histograms of the distribution of FCD values in each matrix are obtained over all participants (blue) and for each simulation (red), and their similarity is evaluated by means of the Kolmogorov-Smirnov distance. (D) Across values of the global coupling parameter *G*, we compute the KS-distance between the empirical FCD and the FCD of each simulation (red solid line; dashed lines indicate standard deviation). The optimum value of *G* for the model (blue vertical line) is chosen as the one that minimises the average KS-distance across 100 simulations (here shown for the awake condition of the propofol dataset, where *G*=1.6). This procedure determines the value that allows the model to best simulate the empirical dynamics of functional connectivity in the healthy human brain.

### Inhibitory modulation from GABA-A receptor distribution reveals a shared mechanism for loss of consciousness

The effects of propofol anaesthesia on the brain were modelled by capitalising on the recently built whole-brain map of GABA-A regional receptor density, generated on the basis of benzodiazepine receptor (BZR) density measured from [^11^C]flumazenil Positron Emission Tomography (PET) ^94^. Incorporating this information in the DMF model allowed us to evaluate the extent to which the dynamics of the anaesthetised brain can be explained in terms of propofol-induced alterations in the detailed balance of local excitation and inhibition.

In previous work Deco and colleagues ^86^ modelled the effects of the serotonergic drug LSD by locally modulating the neuronal gain of each excitatory population in the model according to the empirical distribution of 5HT-2A receptors across brain regions ^86^. Inspired by their approach, here we show for the first time that the influence of regional GABA-A receptor density on functional dynamics can be modelled using a DMF model informed by regional GABA-A receptor density.

The strategy followed in ^86^ was to first calibrate the model on baseline data to obtain a global coupling value, and then fit a secondary excitatory parameter separately on baseline and post-dose data. Our approach follows Deco’s but differs in one key respect: given the inhibitory nature of GABA, we modulated the inhibitory (rather than excitatory) local gain.^1^ To do so, we introduced an inhibitory gain scaling parameter in the model, denoted by *s*_*I*_. This parameter allowed us to scale the inhibitory gain at each region according to the empirical local density of GABA-A receptors, as quantified based on PET-derived maps of receptor density ^94^.

This procedure allowed us to ask whether adjusting the value of inhibitory gain *s*_*I*_ according to local GABA-A receptor density would allow the model to simulate the characteristic dynamics of acute propofol-induced unconsciousness. A positive answer to this question would implicate regional GABA-ergic inhibition as a neurobiological mechanism behind the action of propofol (a known GABA-ergic agonist) in inducing the characteristic macroscale dynamics observed during loss of consciousness due to propofol anaesthesia ^13,38,40,41^.

To address this issue, we studied whether some appropriate value of *s*_*I*_ (which scales the gain related to the local GABA-A receptor density) would improve the model’s ability to simulate the dynamics of deep propofol anaesthesia. For this purpose, we used the previously calibrated DMF model to generate simulations for each value of *s*_*I*_ between 0 (corresponding to the model without local GABA inhibitory modulation) and 1, in increments of 0.02. Then, for each value of *s*_*I*_, we computed the KS distance between the model’s simulated macroscale dynamics and the empirical dynamics observed in the anaesthetised subjects. The optimal value of *s*_*I*_, was then identified as the value that resulted in the minimum mean KS distance between empirical and simulated dynamics (across N=10 simulations for each value of *s*_*I*_), thereby establishing a *propofol model*.

Note that a model incorporating additional neurobiological information (in this case, information about the regional distribution of GABA-A receptors) might produce generally better-fitting simulations just in virtue of its increased complexity. To control for this possibility, we compared the propofol model with an analogous model, which also incorporated regional GABA-A receptor density, but whose inhibitory gain scaling parameter was chosen as the one that best fitted the empirical dynamics observed in awake subjects. We refer to this as the *baseline model*.

Having completed the fitting procedure for our models, we then proceeded to analyse the models’ performance. To this end, we generated 100 simulations from each of the propofol and baseline models. For both models, we then computed the KS distance between each simulation, and the empirical dynamics observed during wakefulness, and during anaesthesia. This provided us with a way to quantify the ability of each model (in terms of goodness of fit, i.e. low KS distance) to simulate the empirical brain dynamics observed during wakefulness, and the empirical brain dynamics observed during propofol-induced loss of consciousness.

Results show that incorporating the regional distribution of GABA-A receptors as local modulators of the global inhibitory gain, substantially improved the DMF model’s fit to empirical brain dynamics observed during propofol anaesthesia (Figure 3B and Table 1). This improvement could not be explained as a generic effect due to additional model complexity: if this had been the case, improvements should have been observed for the model’s ability to fit both awake and anaesthetised dynamics. Instead, the improvement was specific to anaesthetised dynamics, and the propofol model actually had poorer ability to fit the awake dynamics. In other words, the addition of this inhibition parameter in accordance with its empirical distribution across brain regions, makes the model capable of switching between simulating awake or anaesthetised brain dynamics - and when it is simulating anaesthetised brain dynamics, the model’s fit to awake brain dynamics deteriorates. Since propofol is a well-known GABA-ergic agonist, these results confirm that taking into account GABA agonism (local modulation of inhibitory gain by regional GABA-A receptor density) is sufficient to recapitulate the known effects of the GABA-ergic agent propofol on empirical brain dynamics, leading to dynamics that are known to characterise the state of unconsciousness.

**Table 1.**
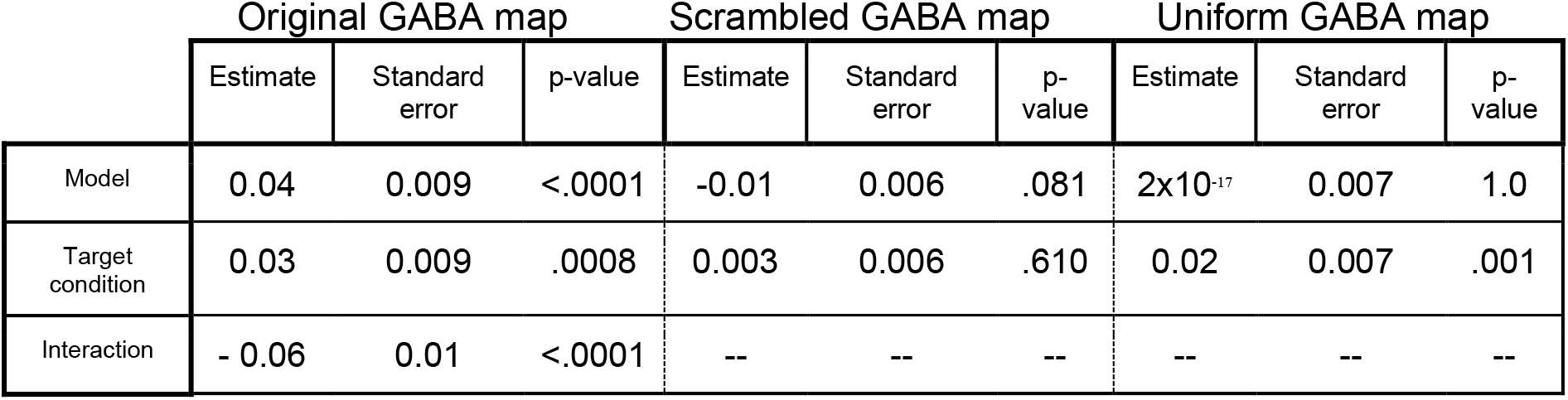
Statistical testing for the effects of local inhibitory GABA modulation.

|  | Original GABA map |  |  | Scrambled GABA map |  |  | Uniform GABA map |  |  |
| --- | --- | --- | --- | --- | --- | --- | --- | --- | --- |
|  | Estimate | Standard error | p-value | Estimate | Standard error | p-value | Estimate | Standard error | p-value |
| Model | 0.04 | 0.009 | <.0001 | -0.01 | 0.006 | .081 | 2x10<sup>-17</sup> | 0.007 | 1.0 |
| Target condition | 0.03 | 0.009 | .0008 | 0.003 | 0.006 | .610 | 0.02 | 0.007 | .001 |
| Interaction | - 0.06 | 0.01 | <.0001 | -- | -- | -- | -- | -- | -- |

**Figure 3.**
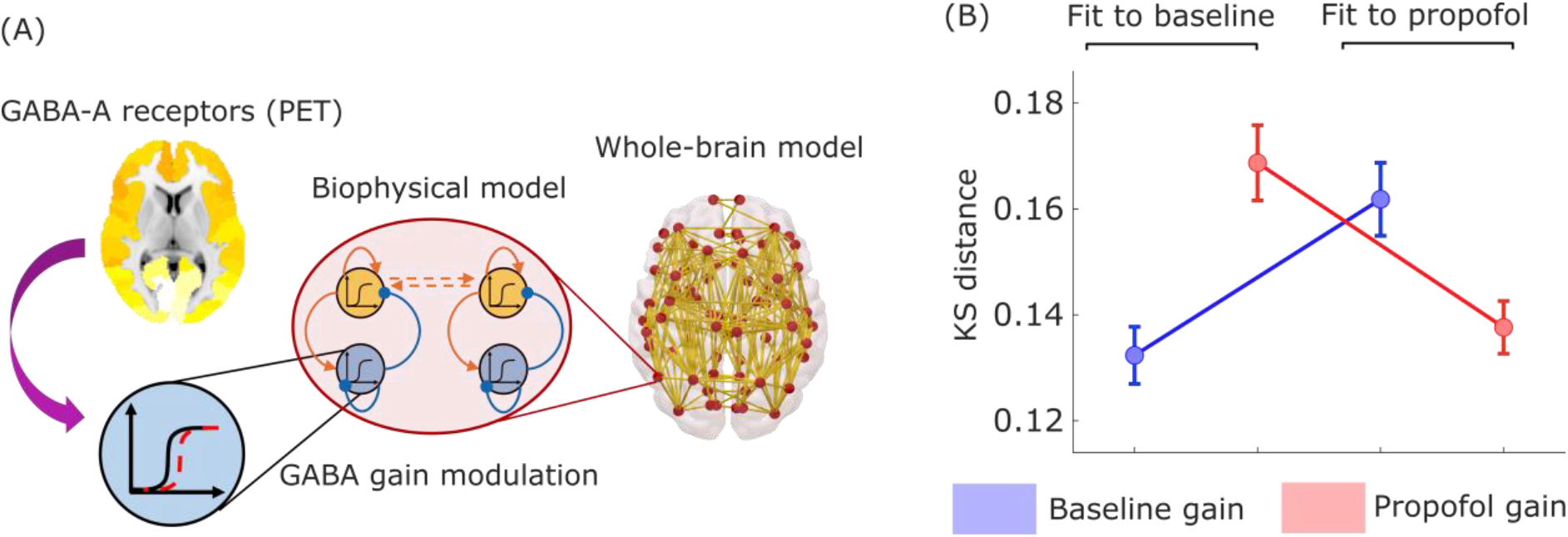
Modulation of inhibitory gain by empirical GABA-A receptor density improves model fit to propofol dynamics. (A) The inhibitory gain of each node in the balanced DMF model is modulated by the regional density of GABA-A receptors, estimated from PET. (B) Mean model fit for 100 simulations for the propofol dataset, quantified as KS-distance to the baseline (awake) and propofol (anaesthetised) conditions, using a value of gain for inhibitory scaling *s*_*I*_ derived from calibrating the model with either the baseline or propofol conditions. Error bars represent the standard error of the mean.

Crucially, we also confirmed that the improved fit to anaesthetised dynamics is not merely the result of increasing overall inhibition in the model: rather, regional information about the distribution of GABA receptor density plays a key role in the model’s improved fit. To demonstrate this point, we show that the results are not replicated if the PET-derived regional distribution of GABA-A receptor density is reshuffled across regions (Figure 4A), or if uniform values are used for each region (i.e. by setting all regions to have a value equal to the mean of the distribution; Figure 4B). In both cases, the model’s ability to fit anaesthetised dynamics is not improved with respect to the baseline model (Figure 4A,B and Table 1). Therefore, our results show that the specific regional distribution of GABA-A receptors across the cortex plays a key role in generating the brain dynamics characteristic of unconsciousness induced by propofol administration.

**Figure 4.**
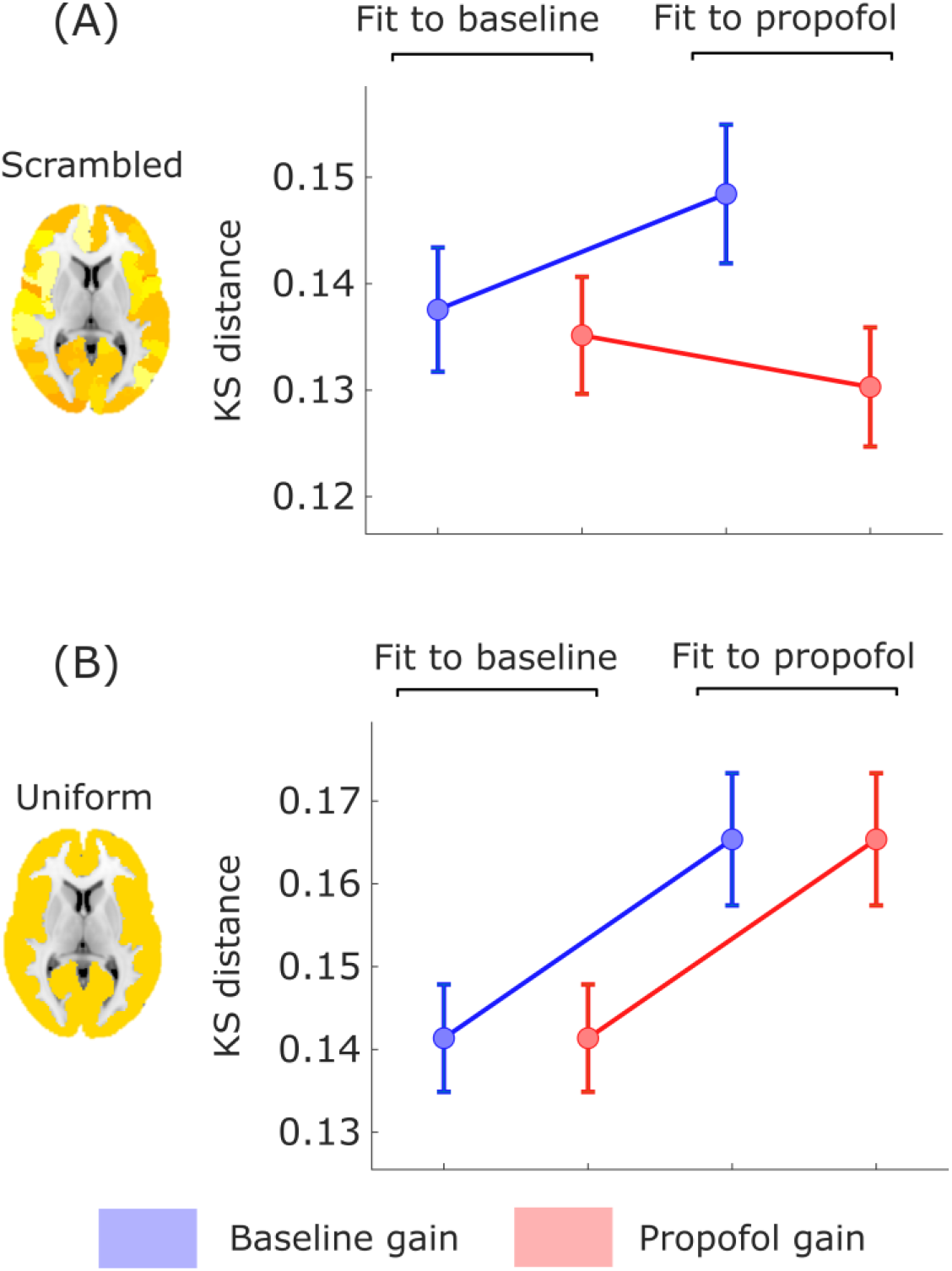
Modulation of inhibitory gain by reshuffled or uniform GABA-A receptor density. (A) The inhibitory gain of each node in the balanced DMF model is modulated by the regional density of GABA-A receptors, reshuffled across cortical regions. Plot shows the mean model fit for 100 simulations, quantified as KS-distance between simulated and empirical dynamics, for each combination of condition and gain. (B) The inhibitory gain of each node in the balanced DMF model is modulated by the regional density of GABA-A receptors, set to a uniform value (mean of the empirical distribution) across cortical regions. Plot shows the mean model fit for 100 simulations, quantified as KS-distance between and empirical dynamics, for each combination of condition and gain. Error bars represent the standard error of the mean.

### Simulated brain injury induces unconscious-like dynamics

Whole-brain computational models provide a unique tool to understand the effects of connectome alterations on macroscale brain dynamics ^70,71,83^. Connectome replacement allows us to determine which of two conditions is more compatible with a given perturbation of the connectome, in terms of the connectome’s capacity to support the corresponding brain dynamics. Specifically, a smaller impact of connectome perturbation on the model’s fit to condition X than condition Y, indicates that the perturbed connectome is better suited to supporting the dynamics of condition X than Y. We term this procedure “*Connectome Replacement Analysis*”, which is illustrated in Figure 5A.

**Figure 5.**
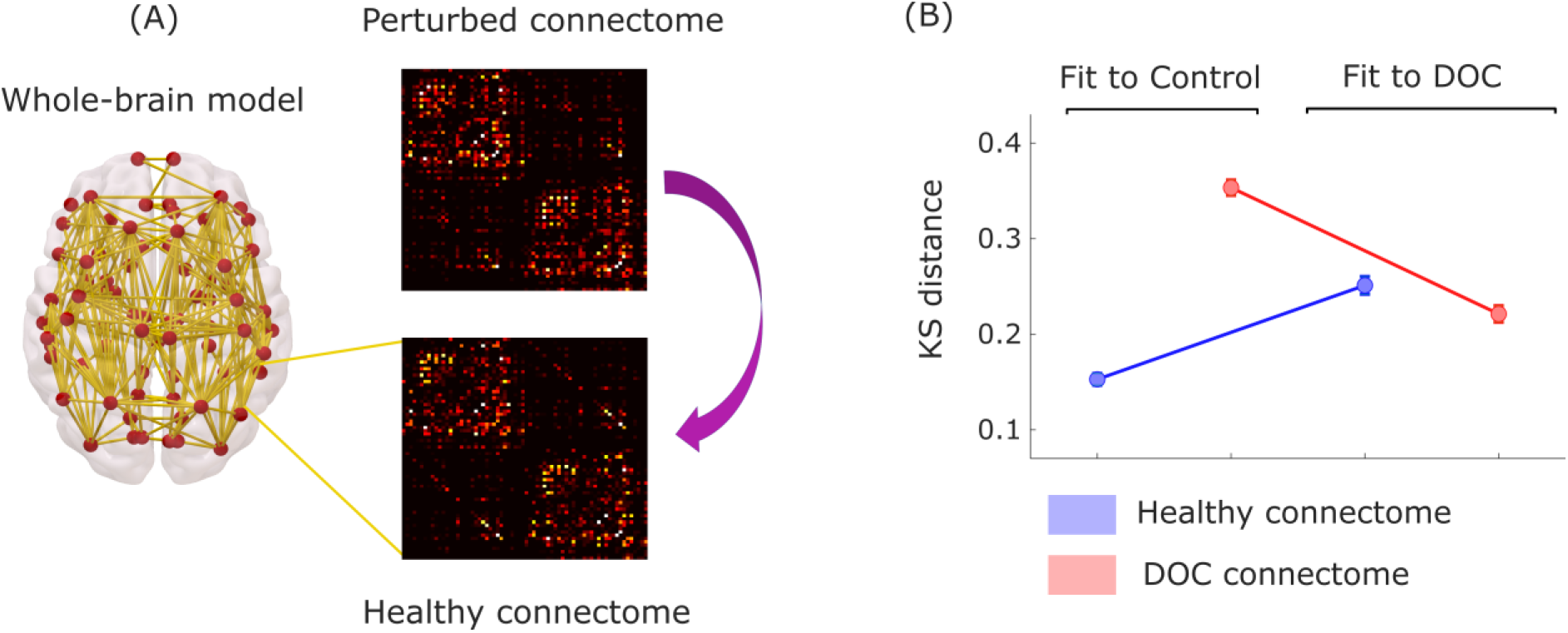
Connectome replacement analysis with DOC connectome. (A) The original healthy connectome of the model is replaced with the group-average connectome obtained from diffusion MRI of N=21 DOC patients, and the resulting model is used to generate 100 simulations. (B) Plot shows the mean model fit for 100 simulations, for the original and perturbed models for the DOC dataset, quantified as the KS-distance to the control (conscious) and DOC (unconscious) conditions. Error bars represent the standard error of the mean.

Leveraging this capability, we subjected the DMF model to the virtual equivalent of severe brain injury: namely, we replaced the underlying connectivity matrix governing the long-range interactions between brain regions with a connectome obtained from diffusion-weighted imaging of N=21 patients with chronic DOC due to severe brain injury. This procedure imparts on the model with effects akin to what severe brain injury does on anatomical connectivity. This “virtual DOC” provides a way to isolate the effects over brain dynamics of connectivity disruptions that result in loss of human consciousness.

Note that such substantial perturbations are expected to deteriorate the model’s ability to replicate the dynamics of awake healthy brains (i.e. increasing the KS distance, corresponding to decreased goodness-of-fit): our models were optimised with biophysical parameters pertaining to healthy brains, and using a healthy connectome. However, our hypothesis was that the dynamics generated by the model with DOC connectome should be more similar (lower KS distance, indicating a better fit) to the empirical dynamics of DOC patients’ brains, than to the dynamics of conscious, healthy brains.

Our results supported these predictions. As expected, connectome replacement led to a reduction in the model’s ability to fit control brain dynamics (Figure 5B). Also as expected, the model with the healthy connectome was better able to simulate conscious than unconscious dynamics (Figure 5B, blue plots). Crucially, however, this pattern reversed following replacement of the healthy connectome with the DOC connectome (Figure 5B, red plots): dynamics generated using the DOC connectome were more similar to the empirical brain dynamics of DOC patients, than to conscious controls (Table 2). This observation supports our hypothesis, demonstrating that unconscious brain dynamics are more compatible with the DOC connectome than conscious dynamics; below, we also demonstrate that this result is not specific to the chronic unconsciousness that characterises disorders of consciousness, but rather it generalises to the transient unconsciousness caused by propofol anaesthesia, too.

**Table 2.** Statistical testing for the effects of connectome replacement.

|  | DOC connectome |  |  | Random connectome |  |  | Lattice connectome |  |  |
| --- | --- | --- | --- | --- | --- | --- | --- | --- | --- |
|  | Estimate | Standard error | p-value | Estimate | Standard error | p-value | Estimate | Standard error | p-value |
| Model | 0.20 | 0.012 | <.0001 | 0.29 | 0.012 | <.0001 | 0.06 | 0.012 | <.0001 |
| Target condition | 0.10 | 0.012 | <.0001 | 0.10 | 0.012 | <.0001 | 0.10 | 0.012 | <.0001 |
| Interaction | - 0.23 | 0.016 | <.0001 | - 0.21 | 0.017 | <.0001 | 0.05 | 0.016 | .007 |

Remarkably, these results could be replicated by replacing the original healthy structural connectome with a randomised version having the same average connectivity ^95^. After perturbation, the model’s fit to DOC brain dynamics became better than the model’s fit to conscious dynamics (Figure 6A and Table 2), suggesting that unconscious dynamics are more compatible than conscious dynamics with a randomised connectome. In contrast, the opposite effect was observed when the original connectome was rewired into a regular (lattice) network, which had a larger impact on the model’s fit to DOC than control brain dynamics (Figure 6B and Table 2). Crucially, the fact that lattice networks further degrade the fit of DOC dynamics demonstrates that not every deviation from the original healthy connectome leads to a higher compatibility with unconscious over conscious dynamics: rather, compatibility depends on the specific topology of the perturbed connectome. Together, these results suggest that DOC dynamics are more compatible with an unstructured connectome.

**Figure 6.**
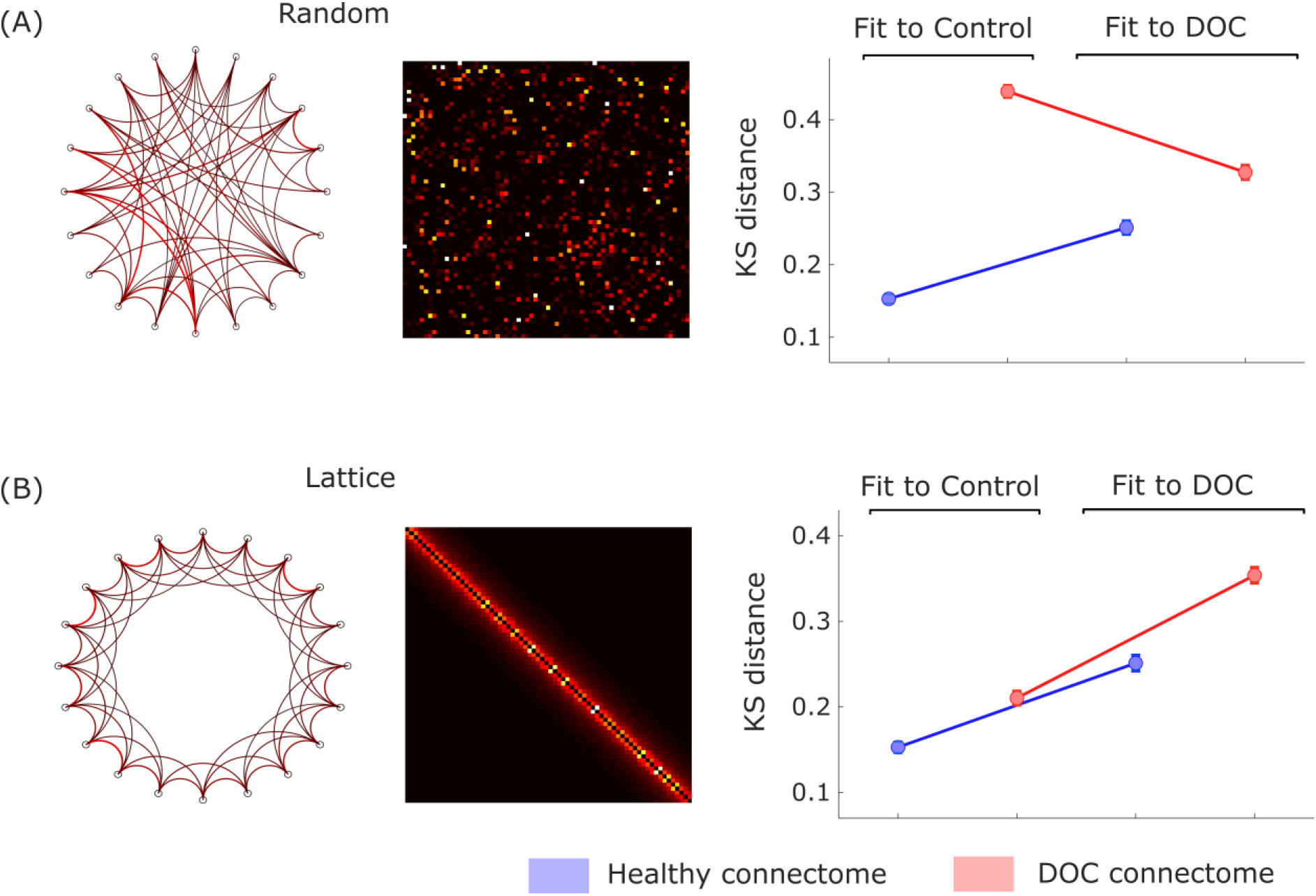
Connectome replacement analysis with random and lattice networks. (A) Plots show the mean model fit for 100 simulations, quantified as the KS distance to each condition for the original and perturbed models using a randomised connectome. (B) Plots show the mean model fit for 100 simulations, quantified as the KS distance to each condition for the original and perturbed models using a lattice-like connectome. Error bars represent the standard error of the mean.

### Generalisation across datasets

Having identified the role of GABA-mediated inhibition for propofol anaesthesia, we next sought to determine to what extent inhibition can also explain the dynamics of unconsciousness arising from severe brain injury. Our rationale was that, even though these patients have not been exposed to GABA-ergic agents but rather owe their condition to severe brain injury, recent evidence suggests similarities of brain dynamics during anaesthesia and disorders of consciousness ^13,38,39,42,96^. A positive answer to this question would further implicate a change in the excitation-inhibition balance, not just in the generation of dynamics pertaining to propofol anaesthesia, but more broadly as a general mechanism responsible for the characteristic dynamics of unconscious states - whether due to anaesthesia or brain injury.

Therefore, we followed the same “virtual anaesthesia” procedure with empirical data from healthy controls and DOC patients. Intriguingly, we observed analogous results: local modulation of inhibitory gain based on GABA-A receptor density allowed the model to substantially improve its fit to DOC patients’ brain dynamics (Figure 7A and Table 3). However, in contrast with propofol anaesthesia, the improvements were also observed when the regional receptor map was scrambled, or replaced by a uniform map (Figure 7B,C and Table 3). Thus, whereas propofol anaesthesia depends on the specific distribution of GABA-A receptors across the cortex, indicating that these receptors are mediating the effects of propofol, the characteristic dynamics of DOCs are less selective, and appear to correspond to a non-specific increase in global inhibition.

**Table 3.** Generalisation of local GABA modulation results to DOC patients.

|  | Original GABA map |  |  | Scrambled GABA map |  |  | Uniform GABA map |  |  |
| --- | --- | --- | --- | --- | --- | --- | --- | --- | --- |
|  | Estimate | Standard error | p-value | Estimate | Standard error | p-value | Estimate | Standard error | p-value |
| Model | 0.08 | 0.011 | <.0001 | 0.01 | 0.012 | <.0001 | 0.07 | 0.011 | <.0001 |
| Target condition | 0.06 | 0.011 | <.0001 | 0.11 | 0.012 | .317 | 0.08 | 0.011 | <.0001 |
| Interaction | - 0.14 | 0.015 | <.0001 | - 0.15 | 0.017 | <.0001 | - 0.18 | 0.016 | <.0001 |

**Figure 7.**
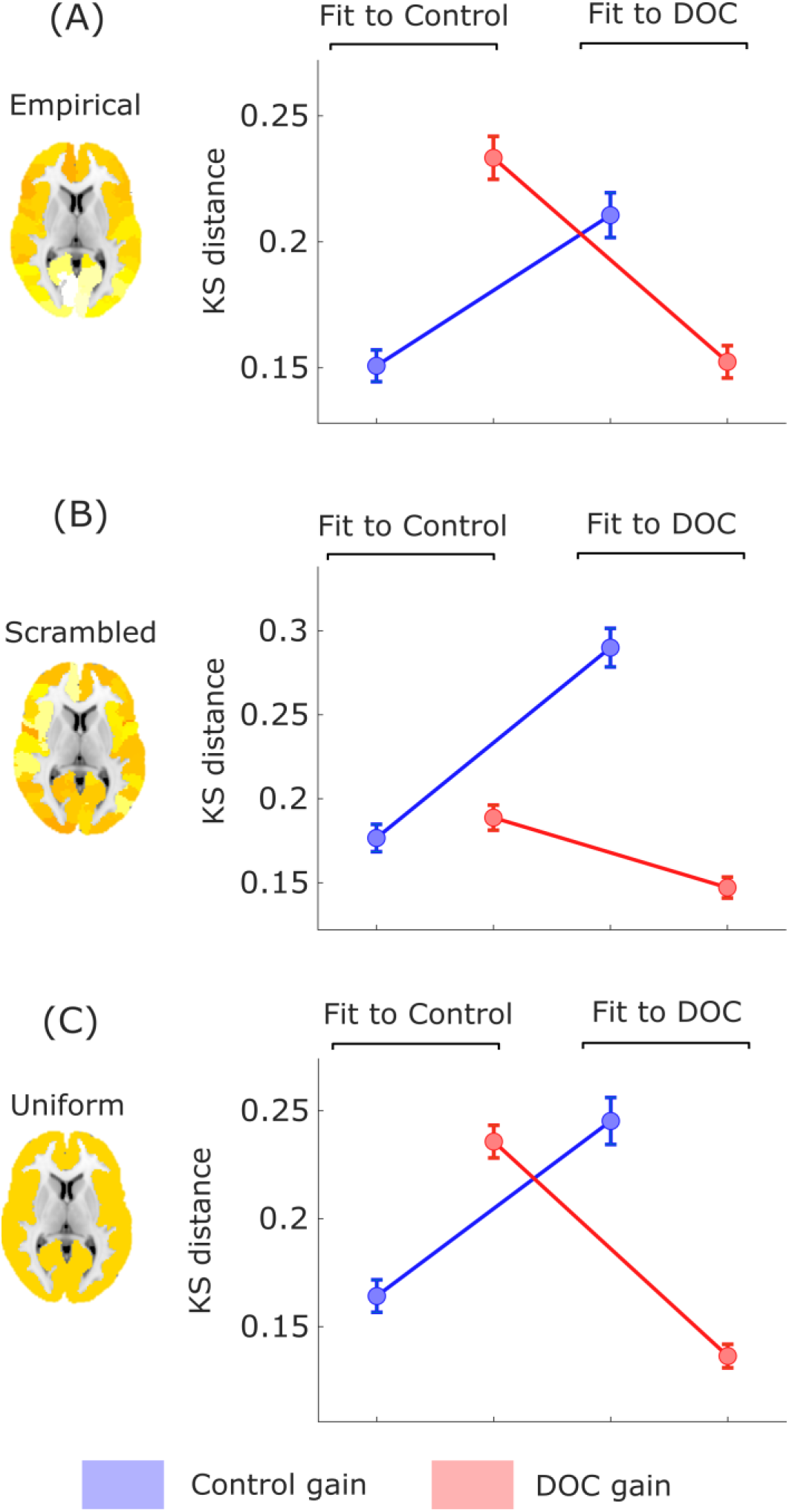
Modulation of inhibitory gain by empirical GABA-A receptor density improves model fit to DOC brain dynamics. (A) Mean model fit for 100 simulations for the DOC dataset, evaluating KS distance (lower means better fit) to the control (conscious) and DOC (unconscious) conditions, using a value of inhibitory gain scaling *s*_*I*_ derived from either the control subjects or the DOC patients. (B) The inhibitory gain of each node in the balanced DMF model is modulated by the regional density of GABA-A receptors, reshuffled across cortical regions. Plot shows the mean model fit (KS-distance) between simulated and empirical dynamics, for each combination of condition and gain. (C) The inhibitory gain of each node in the balanced DMF model is modulated by the regional density of GABA-A receptors, set to a uniform value (mean of the empirical distribution) across cortical regions. Plot shows the mean model fit (KS-distance) between simulated and empirical dynamics, for each combination of condition and gain. Error bars show the standard error of the mean.

Additionally, if propofol and severe injury are different ways by which the human brain can be pushed towards unconsciousness, then inducing a virtual DOC via connectome replacement should also lead to a model that is better able to simulate the dynamics of an anaesthetised brain, than an awake brain - thereby recapitulating what we previously observed with DOC brain dynamics. Remarkably, results show that - as previously observed with DOC patients - simulated dynamics generated from a model using the DOC connectome are more compatible (lower KS distance) with the dynamics of propofol anaesthesia than with the dynamics of awake subjects’ brains (Figure 8A and Table 4).

**Table 4.** Generalisation of connectome replacement results to anaesthetised volunteers.

|  | DOC connectome |  |  | Random connectome |  |  | Lattice connectome |  |  |
| --- | --- | --- | --- | --- | --- | --- | --- | --- | --- |
|  | Estimate | Standard error | p-value | Estimate | Standard error | p-value | Estimate | Standard error | p-value |
| Model | 0.09 | 0.010 | <.0001 | 0.32 | 0.010 | <.0001 | 0.20 | 0.006 | <.0001 |
| Target condition | 0.02 | 0.010 | .031 | 0.02 | 0.010 | .0495 | 0.04 | 0.006 | <.0001 |
| Interaction | - 0.08 | 0.013 | <.0001 | - 0.07 | 0.015 | <.0001 | - | - | - |

**Figure 8.**
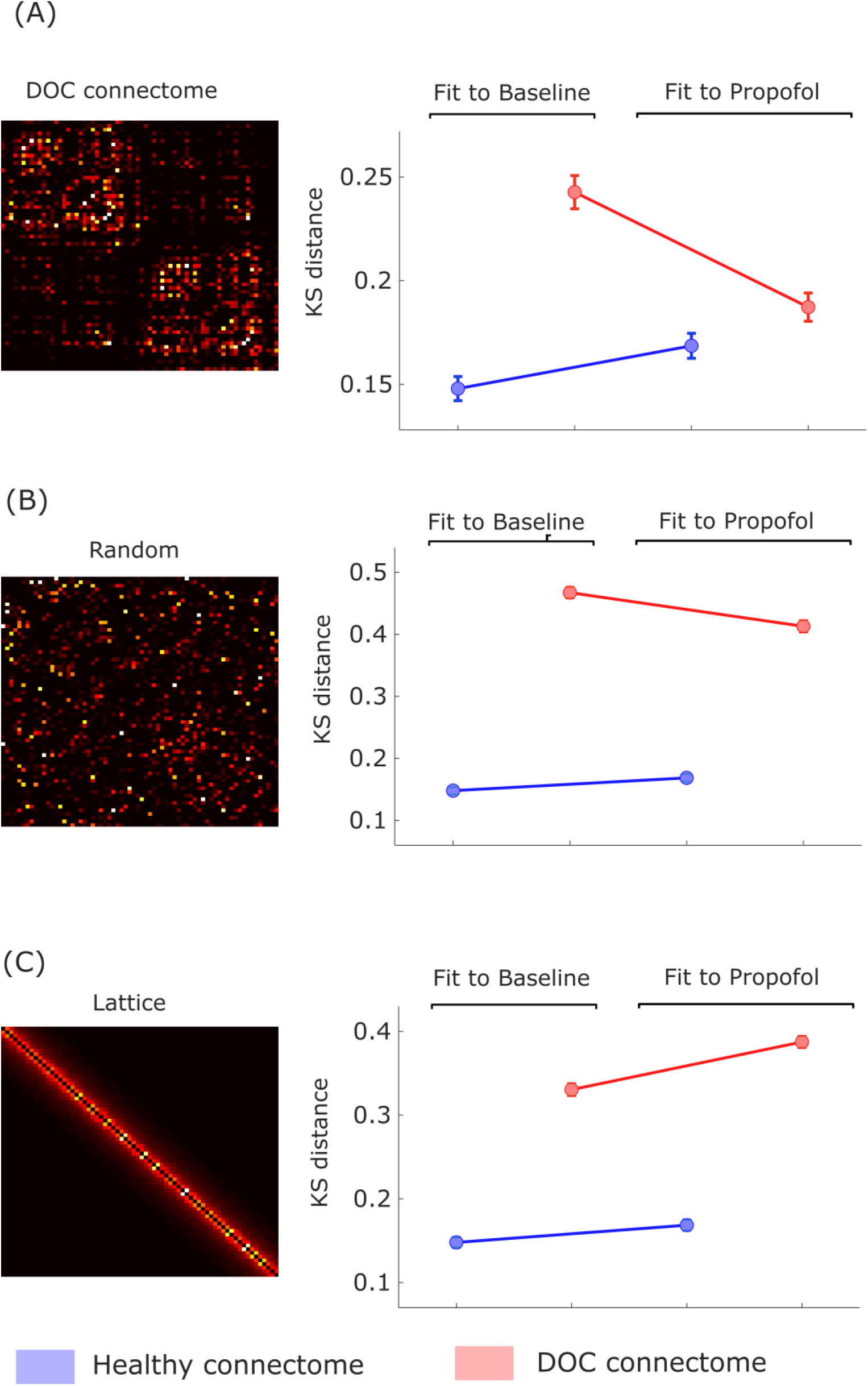
Connectome replacement analysis with DOC connectome generalises to propofol anaesthesia. (A) Plot shows the mean model fit for 100 simulations from the original and perturbed models for the DOC dataset, evaluated as KS-distance to the baseline (conscious) and propofol (unconscious) conditions. (B) Plot shows the mean model fit (KS distance) for 100 simulations, showing the fit to each condition for the original and perturbed models using a randomised connectome. (C) Plot shows the mean model fit (KS distance) for 100 simulations, showing the fit to each condition for the original and perturbed models using a lattice-like connectome. Error bars represent the standard error of the mean.

Furthermore, the importance of topology of the perturbed connectome was also observed for propofol dynamics, with randomisation of the connectome having a smaller effect on the model’s ability to fit propofol dynamics than awake dynamics (Figure 8B and Table 4). Conversely, the decrease in the model’s ability to fit empirical data was exacerbated for the propofol condition, when the original connectome was replaced with a lattice network (Figure 8C and Table 4).

These findings generalise our DOC results to propofol anaesthesia, indicating that the DOC connectome is not only more compatible with DOC patients’ brain dynamics than with healthy subjects’ dynamics. Rather, the generalisation to propofol anaesthesia suggests that the DOC connectome may be more compatible with unconscious dynamics in general: whether arising from brain injury or pharmacological intervention.

## Discussion

The human brain generates a constantly varying set of neural dynamics, supporting a rich repertoire of conscious experiences and cognitive functions ^15–18,20^, which unfold out of an interplay between the structural connectome and the brain’s neuromodulatory mechanisms ^16,31–37^. This paper sought to identify neurobiological mechanisms that are capable of explaining how highly dissimilar causes - such as transient perturbations of neurotransmission *versus* chronic lesions to brain anatomy and connectivity - can give rise to loss of consciousness and its characteristic brain dynamics ^14^.

To this end, we employed a whole-brain *Dynamic Mean Field* model that simulates macroscale functional dynamics of the human brain by means of neurobiologically realistic biophysical modelling, which integrates empirical neuronal dynamics from functional MRI, anatomical connectivity obtained from diffusion MRI, and neurotransmitter receptor density estimated from positron emission tomography. In addition to providing insights about the mechanisms that are responsible for observed macroscale neurobiology, DMF models offer a powerful tool to understand the causal mechanisms underlying pharmacologically-induced macroscale changes in neural dynamics ^86–88^. The effect of inhibition was assessed by enriching the DMF model, modulating the neuronal gain of each inhibitory population according to the empirical density of GABA-A receptors across cortical regions, quantified using in-vivo Positron Emission Tomography autoradiography ^94^. Additionally, the role of lesions to brain anatomy was studied by considering the connectome observed in a population of DOC patients.

Our results demonstrate that GABA-mediated inhibition plays a mechanistic role in the emergence of the characteristic macroscale dynamics observed during propofol-induced unconsciousness. These results align with neurophysiological evidence indicating that propofol is primarily a GABA-A receptor agonist ^46,47^. Indeed, our results further indicate that propofol anaesthesia is crucially dependent on the specific regional distribution of GABA-A receptors across the cortex, since neither reshuffling this distribution across regions nor setting all regions to equal density values could reproduce the same effect. Future research may build on this finding of regional specificity by seeking to identify whether GABA-A receptor density at specific regions plays an especially prominent role in propofol-induced anaesthesia, or whether concerted action across the entire cortex is required. The prominent involvement of the brain’s default mode network in anaesthesia induced with the GABA-ergic agents propofol and sevoflurane ^13,39,40,93,97,98^ and more specifically the precuneus/posterior cingulate cortices, suggests that GABA receptor density at these regions may be an especially promising candidate for predicting anaesthetic effects.

Remarkably, our PET-informed results showed that considering GABA-mediated scaling of regional inhibitory gain also improved the model’s ability to simulate the characteristic dynamics of DOC patients’ brains, even though these patients owe their chronic condition to severe brain injury rather than pharmacological intervention. This observation suggests a change of excitatory-inhibitory balance in favour of inhibition, not just in the generation of dynamics pertaining to propofol anaesthesia, but more broadly as a general neurobiological mechanism for the dynamics that characterise unconsciousness - whether due to anaesthesia or brain injury. Indeed, there is evidence that physiologically awake but unconscious DOC patients show cortical OFF-periods analogous to those observed in healthy individuals during sleep ^99^, possibly arising from reduced cortico-cortical connectivity and a resulting shift in excitatory-inhibitory balance towards excessive inhibition ^100^, as is observed at a local level after stroke ^101^. And indeed, both disorders of consciousness and general anaesthesia are known to correspond to reduced cerebral metabolism, as measured with PET ^102,103^.

Crucially, the present results are consistent with the recent findings of ^41^, where the framework of connectome harmonic decomposition revealed how propofol anaesthesia and disorders of consciousness can both be explained in terms of coarse-grained harmonic modes of the connectome providing increased contributions to brain dynamics. An increase in the contribution of coarse-grained harmonic modes had been previously generated *in silico* by increasing inhibition in a Wilson-Cowan computational model ^16^. Hence, taken together these studies suggest that a change in the global balance between excitation and inhibition in favour of inhibition seems to play a role in the characteristic dynamics of unconsciousness - which the present work formally demonstrates by combining empirical fMRI data on propofol anaesthesia and DOC, with whole-brain computational modelling informed by GABA-A receptor density.

Nevertheless, a key difference emerged between anaesthesia and DOC: whereas anaesthesia critically depends on propofol’s specific pattern of local inhibition across the cortex, incorporating regional specificity of GABA receptor density distribution did not further improve the model’s ability to simulate DOC patients’ brain dynamics, beyond the improvement provided by using a uniform or scrambled GABA-A receptor map. Therefore, whereas propofol anaesthesia is causally mediated by GABA-A receptors and their specific distribution across the cortex, it appears that a global increase in inhibition is sufficient to generate the characteristic dynamics of disorders of consciousness.

Our results from connectome replacement in the whole-brain model informed by empirical anatomical connectivity (dMRI), point to injury-induced randomisation of the connectome as one such candidate mechanism in DOC patients. Specifically, our findings show that (a) unconscious fMRI dynamics (whether due to propofol anaesthesia or brain injury) are more compatible with the empirical DOC connectome, than conscious dynamics; and (b) unconscious dynamics are also more compatible with a random connectome than conscious dynamics (whereas the opposite holds for a lattice-like connectome). It makes sense that the effect of rewiring the network into a lattice should have the opposite effect of randomisation: lattice networks are characterised by high clustering but also high average shortest path between pairs of nodes - the opposite of random networks. Indeed, the well-known “small-world” organisation that characterises many real-world networks, including the human brain ^104,105^ (but see ^106^) is defined as being the topological middle ground between these two extremes of possible network configurations ^95,107^.

It is also remarkable that the same results from connectome replacement - greater compatibility of unconscious dynamics with the DOC connectome perturbation - could be generalised to the propofol dataset. For DOC patients, such a result may perhaps be expected, since the initial connectome used in the model was obtained from healthy controls, whereas the perturbed DOC connectome was obtained by combining the individual connectomes of the same DOC patients. However, observing the same result in the propofol dataset is a powerful validation of our approach, demonstrating that the results are specific to the presence vs absence of consciousness, rather than being influenced by the specific dataset used.

Thus, thanks to connectome replacement we can infer that the increased neuronal inhibition that characterises both disorders of consciousness and anaesthesia, is functionally equivalent to randomisation of the connectome. However, propofol’s anaesthetic effects are mediated by GABA-A receptors according to their specific regional distribution, whereas disorders of consciousness can be explained in terms of a more generic increase in global inhibition - possibly arising from randomisation of the connectome due to anatomical lesions, whose extent and location are do not follow uniform patterns. Anaesthesia may be expected to operate similarly across individuals, in terms of which regions are more or less affected by propofol. In contrast, each DOC patient is unique in the cause, extent and location of their brain injury. As a result, whereas anaesthesia may depend on specific localised patterns, it stands to reason that the characteristic macroscale dynamics of DOC patients’ brains should arise from global-scale neurobiological mechanisms, which may originate from a variety of causes without necessarily depending on specific locations for injury. On the other hand, future research may also explicitly investigate whether lesions leading to disorders of consciousness show an association with regional distribution of GABA-A receptors.

Our results combining fMRI (neural dynamics), dMRI (anatomical connectivity) and PET (neurotransmitter system) demonstrate that human consciousness arises from the delicate balance of local excitation and inhibition, interacting across an intricate network of anatomical connections. Many paths can lead to unconsciousness by disturbing this balance, whether by influencing the nodes’ activity (through inhibitory modulation) or the connectivity between them (through connectome randomisation). As befits such a complex dynamical system as the human brain, it is likely that other paths to unconsciousness will also exist, explaining phenomena such as regular sleep-wake alternation, epileptic seizures, and the effects of non-GABAergic anaesthetics such as ketamine – some of which have already started to be explored using whole-brain computational modelling ^68,76,78,79,81^. Extending the present framework to account for additional ways of losing consciousness will be a crucial endeavour, informing both our understanding of brain network function and of human consciousness. Likewise, it is vital to combine multimodal neuroimaging and whole-brain modelling to identify paths from unconsciousness back to consciousness, using our understanding of post-anaesthetic recovery to restore consciousness in DOC patients, whether by means of custom-designed drugs or deep brain stimulation ^57,58,86^.

### Limitations

A well-known adage asserts that “All models are wrong”, and the present work is no exception. Models of neurobiological function can vary in complexity related to the level of physiological detail and scale, with both aspects incurring costs in terms of computational resources and time. Thus, trade-offs between realism and complexity are unavoidable ^35^. At the end of the spectrum favouring physiological accuracy, microscale spiking models of individual neurons can incorporate thousands of parameters to account for numerous aspects of their neurobiology, but at present such models are not yet able to demonstrate computational scalability and theoretical tractability that could make them useful for the purposes of the present investigation of macroscale brain dynamics ^35^. At the other end of the spectrum, important insights about the spatial and temporal organisation of macroscale brain networks can be attained from abstract statistical modelling based on Hidden Markov Models, PCA/ICA, and clustering approaches ^17,18,20,23,28,29,108^. Rather than seeking to provide plausible neurobiological mechanisms of empirical brain dynamics, these approaches aim to obtain low-dimensional descriptions of the data (in terms of “metastates”, components or clusters), which can lead to conceptually and computationally more tractable accounts that have been proven useful to study questions about the macroscale organisation of the brain.

Note that the same adage also admits that “some models are useful”: following insights from statistical mechanics, which has demonstrated how macroscopic phenomena can sometimes be understood when atoms are modelled as aggregates rather than individuals. The present work uses a dynamic mean-field model to reduce the complexity of spiking neuron models to a more tractable set of differential equations that represent the mean activity of macroscopic neuronal ensembles ^37,90^. This approach allowed us to simulate realistic whole-brain dynamics in terms of a small number of neurobiologically-informed parameters, thus combining tractability with causal insight.

Nevertheless, we acknowledge that a wide variety of other modelling approaches are possible; indeed, even among DMF models, alternatives have been developed that incorporate additional information about regional neurobiology ^109,110^ or use different fitting procedures ^87^. Likewise, alternative models (e.g. Hopf, Ising) have recently been used to investigate loss of consciousness during sleep ^76–81^ anaesthesia ^78–80,82,84,85^ and also disorders of consciousness ^80,82,83^. Though less neurobiologically detailed, such models have been able to provide insights about different aspects of brain function, such as criticality and the predicted effects of applying external perturbations to individual regions. Thus, it is clear that further complementary insights may be obtained by considering additional neurobiological mechanisms and multiple levels of explanation - each of which may require a different modelling approach ^14,35,58^.

We also assumed that the characteristic dynamics of propofol anaesthesia and disorders of consciousness would correspond to loss of consciousness. Although this is a robust finding supported by converging empirical evidence, the assumption may not hold for every state of unconsciousness, and active investigation in the area remains ongoing. Finally, it is important to note that the diversity of disorders of consciousness in terms of aetiology and severity can greatly benefit from an individual-subject approach. Since we considered the cohort of DOC patients as a whole, it is possible that the similar effects of perturbation using a random connectome or the DOC connectome may in fact arise because we obtained a single “DOC connectome” from the combination of several patients, whose individual lesions may be specific but distinct. Likewise, each patient may only exhibit increased inhibition in a specific region, but if such regions differ across patients, then considering them together may result in apparently uniform inhibition. Indeed, analysing such a diverse cohort as a group is always challenging; having demonstrated the efficacy of our modelling approach at the group level, we expect that a fruitful avenue for future research will be to refine our results by considering the specificity of each unique patient condition. In this regard, it is intriguing that some DOC patients can be paradoxically awakened by administration of the drug zolpidem, which is a GABA-ergic agonist ^111^, which suggests that - at least some patients - the causative neurobiological mechanisms may be substantially different from those identified here based on a group-average DOC connectome. Thus, in future efforts we will apply the frameworks developed here to individual patients, to explore their specific deficits and potential avenues to promote recovery at a finer-grained level.

It is also not just the DOC patients who could benefit from an individualised approach: the GABA-A receptor map used here was obtained from an independent sample of volunteers ^94^ and therefore the present work could not take into account individual differences in regional GABA-A receptor density distribution. Such differences may however play an important role in explaining individual susceptibility to anaesthesia with GABA-ergic agents - and potentially predict individual risk of experiencing post-anaesthetic complications, such as emergence delirium ^112,113^. Investigating this possibility will be an important avenue of future research.

### Conclusion

Taken together, our results from a neurobiologically plausible whole-brain computational model demonstrate fundamental similarities, not just between the macroscale brain dynamics that characterise anaesthesia and disorders of consciousness ^13,38–42^ but also between the neurobiological mechanisms from which they can arise - despite the fact that anaesthesia is a transient pharmacological intervention and DOCs are the result of permanent neuroanatomical injury. Both disorders of consciousness and propofol anaesthesia were shown to arise from neurobiological mechanisms that are functionally equivalent to connectome randomisation, and both involve increased perturbed excitation-inhibition balance, as indicated by incorporating into the model information about regional GABA-A receptor density estimated from PET. However, differences also emerged: a global increase in inhibition suffices to explain the macroscale dynamics that characterise disorders of consciousness, whereas the anaesthetic effects of propofol are mediated by GABA-A receptors according to their specific distribution across cortical regions.

Overall, the present findings begin to unravel the neurobiological mechanisms by which different perturbations of the brain’s structure and function - transient pharmacological intervention and chronic neuroanatomical injury - can lead to unconsciousness. Having demonstrated the power of whole-brain computational modelling to address this challenge, the same framework may also prove fruitful to address the reverse problem: namely, how the recovery of consciousness after anaesthesia can inform our ability to restore consciousness in DOC patients.

## Materials and Methods

### Anaesthesia Data

The propofol data employed in this study have been published before ^3,13,114^. For clarity and consistency of reporting, where applicable we use the same wording as our previous study ^13^.

#### Recruitment

As previously reported ^13^, “The propofol data were collected at the Robarts Research Institute in London, Ontario (Canada) between May and November 2014. A total of 19 (18–40 years; 13 males) healthy, right-handed, native English speakers, with no history of neurological disorders were recruited. Each volunteer provided written informed consent, following relevant ethical guidelines, and received monetary compensation for their time. The Health Sciences Research Ethics Board and Psychology Research Ethics Board of Western University (Ontario, Canada) ethically approved this study. Due to equipment malfunction or physiological impediments to anaesthesia in the scanner, data from three participants (1 male) were excluded from analyses, leaving 16” ^13^.

#### Procedure

Resting-state fMRI data were acquired at no sedation (Awake), and Deep sedation (anaesthetised: Ramsay score of 5). As previously reported ^13^: “Ramsay level was independently assessed by two anaesthesiologists and one anaesthesia nurse in the scanning room before fMRI acquisition began, in each condition. Additionally, participants performed two tests: a computerised auditory target-detection task and a memory test of verbal recall, to evaluate their level of wakefulness independently of the assessors. For the Awake condition, participants did not receive a Ramsey score, as this scale is designed for patients in critical care. Instead, they had to be fully awake, alert and communicating appropriately. An infrared camera located inside the scanner was used to monitor wakefulness. For the Deep sedation condition, propofol was administered intravenously using an AS50 auto syringe infusion pump (Baxter Healthcare, Singapore); step-wise sedation increments sedation were achieved using an effect-site/plasma steering algorithm combined with the computer-controlled infusion pump. Further manual adjustments were performed as required to reach target concentrations of propofol, as predicted by the TIVA Trainer (European Society for Intravenous Aneaesthesia, eurosiva.eu) pharmacokinetic simulation program. This software also specified the blood concentrations of propofol, following the Marsh 3-compartment model, which were used as targets for the pharmacokinetic model providing target-controlled infusion. The initial propofol target effect-site concentration was 0.6 *μ*g mL^−1^, with oxygen titrated to maintain SpO2 above 96%. Concentration was then increased by increments of 0.3 *μ*g mL^−1^, and Ramsay score was assessed: if lower than 5, a further increment occurred. Participants were deemed to have reached Ramsay level 5 once they stopped responding to verbal commands, were unable to engage in conversation, and were rousable only to physical stimulation. Data acquisition began once loss of behavioural responsiveness occurred for both tasks, and the three assessors agreed that Ramsay sedation level 5 had been reached. The mean estimated effect-site and plasma propofol concentrations were kept stable by the pharmacokinetic model delivered via the TIVA Trainer infusion pump; the mean estimated effect-site propofol concentration was 2.48 (1.82-3.14) *μ*g mL^−1^, and the mean estimated plasma propofol concentration was 2.68 (1.92-3.44) *μ*g mL^−1^. Mean total mass of propofol administered was 486.58 (373.30-599.86) mg. These values of variability are typical for the pharmacokinetics and pharmacodynamics of propofol. At Ramsay 5 sedation level, participants remained capable of spontaneous cardiovascular function and ventilation. However, since the sedation procedure did not take place in a hospital setting, airway security could not be ensured by intubation during scanning, although two anaesthesiologists closely monitored each participant. Consequently, scanner time was minimised to ensure return to normal breathing following deep sedation. No state changes or movement were noted during the deep sedation scanning for any of the participants included in the study” ^13^.

#### Design

As previously reported ^13^: “In the scanner, subjects were instructed to relax with closed eyes, without falling asleep; 8 minutes of fMRI scan without any task (“resting-state”) were acquired for each participant. Additionally, a separate 5-minute long scan was also acquired while a plot-driven story was presented through headphones to participants, who were instructed to listen while keeping their eyes closed” ^13^. The present analysis focuses on the resting-state data only; the story scan data have been published separately ^84^ and will not be discussed further here.

#### Data Acquisition

As previously reported ^13^: “MRI scanning was performed using a 3-Tesla Siemens Tim Trio scanner (32-channel coil), and 256 functional volumes (echo-planar images, EPI) were collected from each participant, with the following parameters: slices = 33, with 25% inter-slice gap; resolution = 3mm isotropic; TR = 2000ms; TE = 30ms; flip angle = 75 degrees; matrix size = 64×64. The order of acquisition was interleaved, bottom-up. Anatomical scanning was also performed, acquiring a high-resolution T1-weighted volume (32-channel coil, 1mm isotropic voxel size) with a 3D MPRAGE sequence, using the following parameters: TA = 5min, TE = 4.25ms, 240×256 matrix size, 9 degrees FA” ^13^.

#### Disorders of Consciousness Patient Data

The DOC patient functional data employed in this study have been published before ^13,41,115^. For clarity and consistency of reporting, where applicable we use the same wording as our previous study ^13^.

#### Recruitment

As previously reported ^13^: “A sample of 71 DOC patients was included in this study. Patients were recruited from specialised long-term care centres. To be invited to the study, patients must have had a DOC diagnosis, written informed consent to participation from their legal representative, and were capable of being transported to Addenbrooke’s Hospital. The exclusion criteria included any medical condition that made it unsafe for the patient to participate (decision made by clinical personnel blinded to the specific aims of the study) or any reason they are unsuitable to enter the MRI scanner environment (e.g. non-MRI-safe implants), significant pre-existing mental health problems, or insufficient English pre injury. After admission, each patient underwent clinical and neuroimaging testing. Patients spent a total of five days (including arrival and departure days) at Addenbrooke’s Hospital. Coma Recovery Scale-Revised (CRS-R) assessments were recorded at least daily for the five days of admission. If behaviours were indicative of awareness at any time, patients were classified as MCS; otherwise UWS. We assigned MCS- or MCS+ sub-classification if behaviours were consistent throughout the week. The most frequent signs of consciousness in MCS-patients are visual fixation and pursuit, automatic motor reactions (e.g. scratching, pulling the bed sheet) and localisation to noxious stimulation whereas MCS+ patients may, in addition, follow simple commands, intelligibly verbalise or intentionally but inaccurately communicate ^53,54^. Scanning occurred at the Wolfson Brain Imaging Centre, Addenbrooke’s Hospital, between January 2010 and December 2015; medication prescribed to each patient was maintained during scanning. Ethical approval for testing patients was provided by the National Research Ethics Service (National Health Service, UK; LREC reference 99/391). All clinical investigations were conducted in accordance with the Declaration of Helsinki. As a focus of this study was on graph-theoretical properties of the brain, patients were systematically excluded from the final cohort analysed in this study based on the following criteria: 1) large focal brain damage (i.e. more than 1/3 of one hemisphere) as stated by an expert in neuroanatomy blinded to the patients’ diagnoses; 2) excessive head motion during resting state scanning (i.e. greater than 3mm in translation and/or 3 degrees in rotation); 3) suboptimal segmentation and normalization of images; 4) incomplete brain acquisition. A total of 21 adults (13 males; 17-70 years; mean time post injury: 13 months) meeting diagnostic criteria for Unresponsive Wakefulness Syndrome/Vegetative State or Minimally Conscious State due to brain injury were included in this study.” ^13^ (Table 5).

**Table 5:** Demographic information for patients with Disorders of Consciousness.

| Sex | Age | Months post injury | Aetiology | Diagnosis | CRS-R Score | Scan |
| --- | --- | --- | --- | --- | --- | --- |
| **M** | 46 | 23 | TBI | UWS | 6 | 12 dir |
| **M** | 57 | 14 | TBI | MCS- | 12 | 12 dir |
| **M** | 35 | 34 | Anoxic | UWS | 8 | 12 dir |
| **M** | 17 | 17 | Anoxic | UWS | 8 | 12 dir |
| **F** | 31 | 9 | Anoxic | MCS- | 10 | 12 dir |
| **F** | 38 | 13 | TBI | MCS | 11 | 12 dir |
| **M** | 29 | 68 | TBI | MCS | 10 | 63 dir |
| **M** | 23 | 4 | TBI | MCS | 7 | 63 dir |
| **F** | 70 | 11 | Cerebral bleed | MCS | 9 | 63 dir |
| **F** | 30 | 6 | Anoxic | MCS- | 9 | 63<br>dir |
| **F** | 36 | 6 | Anoxic | UWS | 8 | 63<br>dir |
| **M** | 22 | 5 | Anoxic | UWS | 7 | 63<br>dir |
| **M** | 40 | 14 | Anoxic | UWS | 7 | 63<br>dir |
| **F** | 62 | 7 | Anoxic | UWS | 7 | 63<br>dir |
| **M** | 46 | 10 | Anoxic | UWS | 5 | 63<br>dir |
| **M** | 21 | 7 | TBI | MCS | 11 | 63<br>dir |
| **M** | 67 | 14 | TBI | MCS- | 11 | 63<br>dir |
| **F** | 55 | 6 | Hypoxia | UWS | 12 | 63<br>dir |
| **M** | 28 | 14 | TBI | MCS | 8 | 63<br>dir |
| **M** | 22 | 12 | TBI | MCS | 10 | 63<br>dir |
| **F** | 28 | 8 | ADEM | UWS | 6 | 63<br>dir |

CRS-R, Coma Recovery Scale-Revised; UWS, Unresponsive Wakefulness Syndrome; MCS, Minimally Conscious State; TBI, Traumatic Brain Injury.

#### FMRI Data Acquisition

As previously reported ^13^: “Resting-state fMRI was acquired for 10 minutes (300 volumes, TR=2000ms) using a Siemens Trio 3T scanner (Erlangen, Germany). Functional images (32 slices) were acquired using an echo planar sequence, with the following parameters: 3 x 3 x 3.75mm resolution, TR = 2000ms, TE = 30ms, 78 degrees FA. Anatomical scanning was also performed, acquiring high-resolution T1-weighted images with an MPRAGE sequence, using the following parameters: TR = 2300ms, TE = 2.47ms, 150 slices, resolution 1 x 1 x 1mm”.

#### Acquisition of Diffusion-Weighted Data

As the data were acquired over the course of several years, two different diffusion-weighted image acquisition schemes were used for the DOC patients. The first acquisition scheme (used for the N=7 patients whose data were acquired earliest in time) used an echo planar sequence (TR = 8300 ms, TE = 98 ms, matrix size = 96 x 96, 63 slices, slice thickness = 2 mm, no gap, flip angle = 90°). This included diffusion sensitising gradients applied along 12 non-collinear directions with 5 b-values that ranged from 340 to 1590 s/mm2 and 5 b = 0 images. One of these patients only had partial brain coverage for the diffusion MRI acquisition, and their DWI data were excluded from further analysis. The more recent acquisition scheme (used for the more recently scanned patients, and for all healthy controls) instead involved the use of 63 directions with a b-value of 1000 s/mm2. Both DWI acquisition types have been used before in the context of structural connectivity analysis in DOC patients ^51,53^.

### Healthy controls

We also acquired diffusion MRI data from N=20 healthy volunteers (13 males; 19-57 years), with no history of psychiatric or neurological disorders. The mean age was not significantly different between healthy controls (M = 35.75; SD = 11.42) and DOC patients (M = 38.24; SD = 15.96) (*t*(39) = −0.57, *p* = 0.571, Hedges’s *g* = −0.18; permutation-based t-test).

#### FMRI Data Acquisition

Resting-state fMRI was acquired for 5:20 minutes (160 volumes, TR=2000ms) using a Siemens Trio 3T scanner (Erlangen, Germany). The acquisition parameters were the same as those for the DOC patients: Functional images (32 slices) were acquired using an echo planar sequence, with the following parameters: 3 x 3 x 3.75mm resolution, TR = 2000ms, TE = 30ms, 78 degrees FA. High-resolution T1-weighted anatomical images were also acquired, using an MPRAGE sequence with the following parameters: TR = 2300ms, TE = 2.47ms, 150 slices, resolution 1 x 1 x 1mm. Two subjects had incomplete fMRI acquisition, leaving N=18 subjects for the final analysis.

#### Acquisition of diffusion-weighted imaging data

The diffusion-weighted acquisition scheme was the same 63-directions scheme used for the DOC patients: TR = 8300 ms, TE = 98 ms, matrix size = 96 x 96, 63 slices, slice thickness = 2 mm, no gap, flip angle = 90°, 63 directions with a b-value of 1000 s/mm2.

### Functional MRI preprocessing and denoising

Following our previous work ^13^, we preprocessed the functional imaging data using a standard pipeline, implemented within the SPM12-based (http://www.fil.ion.ucl.ac.uk/spm) toolbox CONN (http://www.nitrc.org/projects/conn), version 17f ^116^. As described, “The pipeline comprised the following steps: removal of the first five scans, to allow magnetisation to reach steady state; functional realignment and motion correction; slice-timing correction to account for differences in time of acquisition between slices; identification of outlier scans for subsequent regression by means of the quality assurance/artifact rejection software *Artifact Detection Toolbox (art;* (http://www.nitrc.org/projects/artifact_detect); spatial normalisation to Montreal Neurological Institute (MNI-152) standard space with 2mm isotropic resampling resolution, using the segmented grey matter image from each volunteer’s high-resolution T1-weighted image, together with an *a priori* grey matter template” ^13^.

For the DOC patients, due to the presence of deformations caused by brain injury, rather than relying on automated pipelines, each patient’s brain was individually preprocessed using SPM12, with visual inspections after each step. To further reduce potential movement artifacts, data underwent despiking with a hyperbolic tangent squashing function. Since the controls had a shorter scan duration than DOC patients, to ensure comparability between the two cohorts the DOC functional scans were truncated to be of the same length as the control subjects’ functional scans (after removal of the initial scans).

To reduce noise due to cardiac and motion artifacts, we applied the anatomical CompCor method of denoising the functional data ^117^. The anatomical CompCor method (also implemented within the CONN toolbox) involves regressing out of the functional data the following confounding effects: the first five principal components attributable to each individual’s white matter signal, and the first five components attributable to individual cerebrospinal fluid (CSF) signal; six subject-specific realignment parameters (three translations and three rotations) as well as their first-order temporal derivatives; the artifacts identified by *art;* and main effect of scanning condition ^117^. Linear detrending was also applied, and the subject-specific denoised BOLD signal timeseries were band-pass filtered to eliminate both low-frequency drift effects and high-frequency noise, thus retaining frequencies between 0.008 and 0.09 Hz.

### DWI Preprocessing and tractography

The diffusion data were preprocessed with MRtrix3 tools ^118^. After manually removing diffusion-weighted volumes with substantial distortion ^53^, the pipeline involved the following steps: (i) DWI data denoising by exploiting data redundancy in the PCA domain ^119^ (*dwidenoise* command); (ii) Correction for distortions induced by eddy currents and subject motion by registering all DWIs to b0, using FSL’s *eddy* tool (through MRtrix3 *dwipreproc* command); (iii) rotation of the diffusion gradient vectors to account for subject motion estimated by *eddy* ^120^; (iv) b1 field inhomogeneity correction for DWI volumes (*dwibiascorrect* command); (v) generation of a brain mask through a combination of MRtrix3 *dwi2mask* and FSL *BET* commands.

DTI data were reconstructed from the preprocessed DWIs using DSI Studio (www.dsi-studio.labsolver.org), which implements q-space diffeomorphic reconstruction (QSDR ^121^), an established methodology to investigate structural networks in DOC patients ^54^. As explained in the original publication ^121^, “QSDR is a model-free method that calculates the orientational distribution of the density of diffusing water in a standard space, to conserve the diffusible spins and preserve the continuity of fiber geometry for fiber tracking. QSDR first reconstructs diffusion-weighted images in native space and computes the quantitative anisotropy (QA) in each voxel ^121^. These QA values are used to warp the brain to a template QA volume in Montreal Neurological Institute (MNI) space using the statistical parametric mapping (SPM) nonlinear registration algorithm. Once in MNI space, spin density functions (SDFs) were again reconstructed with a mean diffusion distance of 1.25 mm using three fiber orientations per voxel” ^121^.

Following previous work ^122^, “After reconstruction with QSDR, we used deterministic fiber tracking with a high-performing “FACT” algorithm with 1,000,000 streamlines, to identify the connections between brain regions, following previously established parameters ^122,123^: angular cutoff = 55°, step size = 1.0 mm, tract length between 10mm (minimum) and 400mm (maximum), no spin density function smoothing, and QA threshold determined by DWI signal in the cerebro-spinal fluid. DSI Studio automatically applies a default anisotropy threshold of 0.6 Otsu’s threshold to the anisotropy values of the spin density function, in order to generate a white matter mask which is then used for automatic screening of each streamline, to exclude streamlines improper termination locations ^122,123^”.

### Brain parcellation

For both BOLD and DWI data, brains were parcellated into 68 cortical regions of interest (ROIs), according to the Desikan-Killiany anatomical atlas ^124^, in line with previous whole-brain modelling work ^109^.

### Functional Connectivity Dynamics

Following ^86^, functional connectivity dynamics (FCD) were quantified in terms of Pearson correlation between regional BOLD timeseries, computed within a sliding window of 30 TRs with increments of 3 TRs. Subsequently, the resulting matrices of functional connectivity at times t_x_ and t_y_ were themselves correlated, for each pair of timepoints t_x_ and t_y_, thereby obtaining an FCD matrix of time-versus-time correlations. Thus, each entry in the FCD matrix represents the similarity between functional connectivity patterns at different points in time.

### Group Structural Connectivity

The structural connectivity (SC) for the DMF model was obtained by following the procedure described in ^109^ to derive a group-consensus structural connectivity matrix. Separately for the healthy controls and DOC patients, a consensus matrix *C* was obtained as follows. For each pair of regions *i* and *j*, if more than half of subjects had non-zero connection *i* and *j*, *C*_*ij*_ was set to the average across all subjects with non-zero connections between *i* and *j*. Otherwise, *C*_*ij*_ was set to zero.

### Whole-brain computational modelling

Whole-brain spontaneous brain activity (as quantified using blood oxygen level dependent (BOLD) signal data from functional MRI) was simulated using a neurobiologically realistic Dynamic Mean Field (DMF) model. The DMF model ^31,34,37^ uses an empirically validated mathematical mean-field approach to represent the collective behaviour of integrate-and-fire neurons by means of coupled differential equations, providing a neurobiologically plausible account of regional neuronal firing rate.

Specifically, the model simulates local biophysical dynamics of excitatory (NMDA) and inhibitory (GABA) neuronal populations, interacting over long-range neuroanatomical connections (white matter tracts obtained from diffusion MRI). The model further incorporates multimodal neuroimaging information about empirical brain dynamics (measured using functional MRI) and neurotransmitter receptor density, estimated from positron emission tomography (PET) ^86^.

Each cortical area *n* (defined by a parcellation scheme) is represented in terms of two reciprocally coupled neuronal masses, one excitatory and the other inhibitory, with the synaptic connections between excitatory neuronal populations in different regions given by the weight of structural connectivity, to account for the number and density of interregional axon fibers. Additional factors that can influence the long-range excitatory-to-excitatory coupling between brain regions, such as neurotransmission but also synaptic plasticity mechanisms, are accounted for by a global coupling parameter, *G*. Since conductivity of the white matter fibers is assumed to be constant across the brain, *G* constitutes the only free parameter in the model.

The following differential equations therefore govern the model’s behaviour:

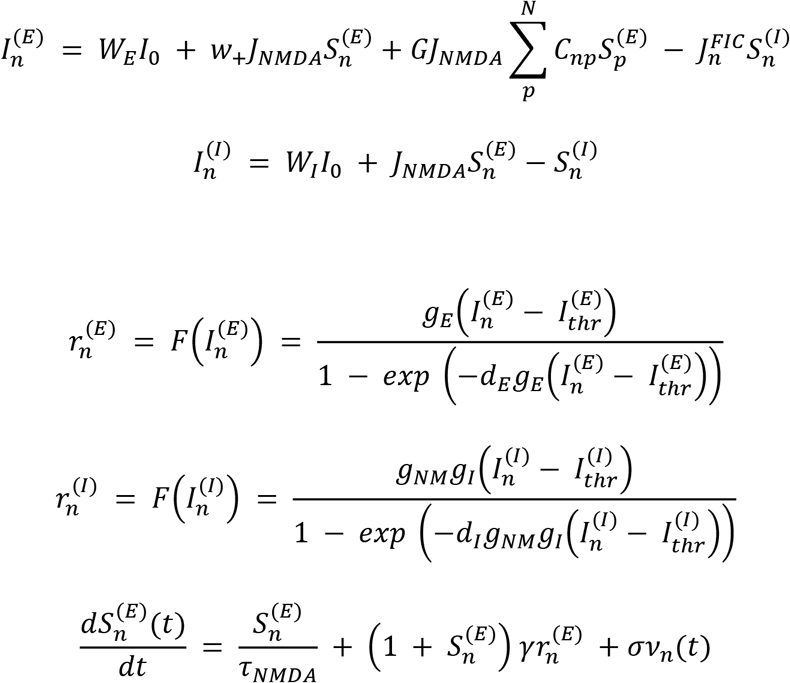

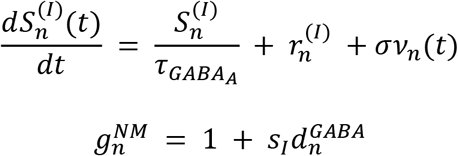

Following previous work ^86,88^, “for each excitatory *(E)* and inhibitory *(I)* neural mass, the quantities 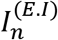, 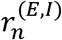, and 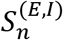 represent its total input current (nA), firing rate (Hz) and synaptic gating variable, respectively. The function F(·) is the transfer function (or *F–I curve*), representing the non-linear relationship between the input current and the output firing rate of a neural population. Finally, 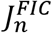 is the local feedback inhibitory control of region *n*, which is optimized to keep its average firing rate at approximately 3Hz ^37,88^, and *v*_*n*_ is uncorrelated Gaussian noise injected to region *n*”. The model’s fixed parameters are reported in Table 6 ^37,86,88^. Additionally, 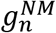 is the *neuromodulatory scaling factor* modulating the transfer function for each cortical region in the model as a function of 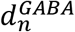, the regional density of GABA-A receptors (see below for details) and an inhibitory gain scaling parameter *s*_*I*_. The *baseline* model (corresponding to a DMF model without GABA-A regional inhibitory neuromodulation) is obtained by setting *s*_*I*_ to zero, in which case *G* remains the sole free parameter in the model. Details for optimisation of the *s*_*I*_ parameter for the GABA-A modulated model are provided below.

**Table 6.** Dynamic Mean Field model parameters from ^37,86,88^.

| Parameter | Symbol | Value |
| --- | --- | --- |
| External current | $I_0$ | 0.382 nA |
| Excitatory scaling factor for $I_0$ | $W_E$ | 1 |
| Inhibitory scaling factor for $I_0$ | $W_I$ | 0.7 |
| Local excitatory recurrence | $w_+$ | 1.4 |
| Excitatory synaptic coupling | $J_{NMDA}$ | 0.15 nA |
| Threshold for $F(I_n^{(E)})$ | $I_{thr}^{(E)}$ | 0.403 nA |
| Threshold for $F(I_n^{(I)})$ | $I_{thr}^{(I)}$ | 0.288 nA |
| Gain factor of $F(I_n^{(E)})$ | $g_E$ | 310 nC 1 |
| Gain factor of $F(I_n^{(I)})$ | $g_I$ | 615 nC 1 |
| Shape of $F(I_n^{(E)})$ around $I_{thr}^{(E)}$ | $dE$ | 0.16 s |
| Shape of $F(I_n^{(I)})$ around $I_{thr}^{(I)}$ | $dI$ | 0.087 s |
| Excitatory kinetic parameter | $\gamma$ | 0.641 |
| Amplitude of uncorrelated Gaussian noise $v_n$ | $\sigma$ | 0.01 nA |
| Time constant of NMDA | $\tau_{NMDA}$ | 100 ms |
| Time constant of GABA | $\tau_{GABA}$ | 10 ms |

A Balloon-Windkessel (BW) hemodynamic model ^92^ was then used to turn simulated regional neuronal activity into simulated regional BOLD signal. The Balloon-Windkessel model considers the BOLD signal as a nonlinear function of the normalized total deoxyhemoglobin voxel content, normalized venous volume, resting net oxygen extraction fraction by the capillary bed, and resting blood volume fraction. The BOLD-signal estimation for each brain area is computed from the level of neuronal activity in that particular area. Finally, simulated regional BOLD signal was bandpass filtered in the same range as the empirical data (0.008-0.09 Hz).

#### Implementation

The code used to run all the simulations in this study was written in optimised C++ using the high-performance library [Eigen](https://eigen.tuxfamily.org/index.php?). The C++ core of the code, together with Python and Octave/Matlab interfaces is publicly available at http://www.gitlab.com/concog/fastdmf.

To simulate BOLD data, FastDMF splits the problem in two steps: integrating the coupled differential equations underlying the DMF model, to obtain excitatory firing rates in each brain region; and using these firing rates to integrate the (uncoupled) differential equations of the BW hemodynamic model and obtain BOLD timeseries.

Integration of the DMF equations is performed with the Euler-Maruyama method, and it is highly parallelizable and bounded by the O(N^2) complexity of the matrix-vector multiplication corresponding to the excitatory-to-excitatory coupling between brain regions.

Simulated excitatory firing rates are stored in a cylindrical array with a fixed buffer size to limit memory requirements.

In addition, a further set of threads is spawned to solve the BW model using the simulated excitatory firing rates. Since the BW solver reads from the same cylindrical array, it interfaces with the DMF solver with a controlled multi-threaded architecture. Every TR-equivalent in simulation time the value of all BOLD signals is copied to a pre-allocated array, to be returned at the end of the requested simulation time.

In a standard laptop, FastDMF attains a speed-up of between 5x and 10x over publicly available Matlab implementations, due to the speed of Eigen and the parallelisation of DMF and BW solvers. In addition, due to the cylindrical buffer, this implementation is able to simulate arbitrarily long BOLD time series with a fixed memory overhead, thereby allocating orders of magnitude less memory than a naive Matlab implementation.

Finally, the library includes interface functions for Matlab (via its [C Matrix API](https://www.mathworks.com/help/matlab/cc-mx-matrix-library.html)) and Python (via the [Boost.Python](https://www.boost.org/) library). In both languages the function returns a standard array (numpy.ndarray in the case of Python) that can be easily processed for further analysis.

#### Fitting of the G parameter

In order to identify appropriate parameters for the simulations, early whole-brain modelling efforts used the grand average FC as target for fitting the model to empirical data. However, it has since become apparent that the macroscale neural signals measured by functional MRI are not static, even on the timescale of a few tens of seconds: rather, they exhibit a wide range of dynamics. Therefore, in order to properly take into account the time-dependencies of FC, it is advantageous to fit the model to empirical functional connectivity dynamics (FCD). Doing so ensures that the simulated BOLD data will exhibit realistic patterns of time-evolving functional connectivity ^86,91^.

Unlike matrices of inter-regional connectivity, where each brain region is the same across different scans, FCDs are represented as matrices encoding the relationship between brain dynamics at different timepoints. Since timepoints are not the same across individuals or scans, FCD matrices cannot be compared by means of simple correlation. Therefore, to evaluate model performance in terms of producing meaningful temporal dynamics, here we follow the approach of ^86^, using the Kolmogorov-Smirnov distance to compare the histograms of empirical and simulated FCD values (obtained from the upper triangular FCD matrix), to find the G parameter that results in the best match between empirical and simulated functional connectivity dynamics.

To find the value of *G* that generates simulations whose FCD best match empirical FCD, we generated 100 simulations for each value of *G* between 0.1 and 2.5, using increments of 0.1. For each simulation at each value of G, we computed the KS distance between empirical (group-wise) and simulated FCD. Finally, we set the model’s G parameter to the value that minimised the mean KS distance - corresponding to the model that is best capable of simulating the temporal dynamics of functional connectivity observed in the healthy human brain at rest.

This procedure was performed separately for the propofol dataset (with 250 TRs) and the DOC dataset, which was truncated to the number of TRs available for the healthy controls (155 TRs).

### Local inhibitory gain modulation from GABA-A maps

We modulated local inhibitory gain based on the recent high-resolution quantitative atlas of human brain GABA-A receptors, generated on the basis of benzodiazepine receptor (BZR) density measured from [^11^C]flumazenil Positron Emission Tomography (PET) autoradiography ^94^. Based on this atlas, we obtained a quantitative measure of GABA-A receptor density for each region of the Desikan-Killiany cortical parcellation. Following ^86^ regional density values were normalised between 0 and 1 by dividing each by the maximum value.

We then used the previously calibrated DMF model to generate simulations for each value of *s*_*I*_ between 0 (corresponding to the model without local GABA-A inhibitory modulation, i.e. the original DMF model) and 1, in increments of 0.02. Then, for each value of *s*_*I*_, we computed the KS distance between the model’s simulated macroscale dynamics and the empirical dynamics observed in each condition (baseline or propofol, control or DOC). For each condition, the optimal value of *s*_*I*_, was then identified as the value that resulted in the minimum mean KS distance between empirical and simulated dynamics (across N=10 simulations for each value of *s*_*I*_).

As validation analysis, we also repeated the same procedure, optimising the inhibitory gain scaling *s*_*I*_, but with two different kinds of receptor density maps: a “scrambled” map, whereby the values of GABA-A receptor density obtained from PET were randomised across regions; and a “uniform” map, whereby each region was set to the same value, corresponding to the mean of the distribution of PET-derived receptor densities.

### Connectome Replacement

Connectome replacement was performed using the initial balanced DMF model (i.e. with optimised *G* parameter, but without additional inhibitory gain modulation), based on the consensus connectome from diffusion imaging of healthy controls (referred to as the “healthy connectome”).

Three perturbed connectomes were used. Firstly, the consensus connectome obtained from diffusion imaging of N=21 DOC patients, referred to as the “DOC connectome”. Secondly, the original healthy connectome was randomised according to the weight-preserving procedure of ^95^ to generate a “random connectome” that differs from the original in terms of topology, but preserves the weight distribution. Thirdly, we used the procedure described in ^95^ to turn the healthy connectome into a lattice network with the same weight distribution - providing a different and opposite perturbation of the network’s topology.

For each perturbed connectome, the DMF model was used to generate 100 simulations, using the optimal global coupling *G*, but with inter-regional connectivity given by the perturbed connectome rather than the original connectome. This was repeated for each dataset (propofol and DOC) and the resulting simulations were compared with each condition (baseline/propofol and control/DOC) in terms of KS-distance.

### Statistical analysis

To test the effect of the various procedures over the DMF model, we performed linear regression modelling using the KS distance as target variable, while considering model and target condition as predictor variables. For each scenario, models with and without interactions between model and target condition were constructed; then, selection between these models was conducted according to the Akaike information criterion (AIC). All analyses were conducted using RStudio (Version 1.3.1093; http://www.rstudio.com/).

## Acknowledgements

The authors would like to thank all the participants for their contribution to this study. This work was supported by grants from the UK Medical Research Council [U.1055.01.002.00001.01 to AMO and JDP]; The James S. McDonnell Foundation [to AMO and JDP]; and the Canada Excellence Research Chairs program (215063 to AMO); the National Institute for Health Research (NIHR, UK), Cambridge Biomedical Research Centre and NIHR Senior Investigator Awards [to DKM], the Stephen Erskine Fellowship (Queens’ College, Cambridge, to EAS), the L’Oreal-Unesco for Women in Science Excellence Research Fellowship to LN; the British Oxygen Professorship of the Royal College of Anaesthetists [to DKM] the Gates Cambridge Trust (to AIL) and the Vice-Chancellor Award (to PC). PAM and DB are funded by the Wellcome Trust (grant no. 210920/Z/18/Z). FR is funded by the Ad Astra Chandaria foundation. The research was also supported by the NIHR Brain Injury Healthcare Technology Co-operative based at Cambridge University Hospitals NHS Foundation Trust and University of Cambridge. AMO and DKM are Fellows of the CIFAR Brain, Mind, and Consciousness Programme.

## Author Contributions

AIL: conceived the study; analysed data; wrote first draft of the manuscript. PAM: conceived the study; contributed to data analysis and interpretation of results; reviewed and edited the manuscript. FR: conceived the study; contributed to data analysis and interpretation of results; reviewed and edited the manuscript. M.M.C., P.C., A.R.D.P: contributed to data analysis. DKM: reviewed the manuscript. DB: reviewed and edited the manuscript. EAS: conceived the study; reviewed and edited the manuscript. P.F., G.B.W., J.A., J.D.P., A.M.O., L.N., D.K.M. and E.A.S. were involved in designing the original studies for which the present data were collected. P.F., M.M.C., G.B.W., J.A., L.N. and E.A.S. all participated in data collection.

## Competing Interests

The authors declare no competing interests.

## Data and Code Availability

The CONN toolbox is freely available online (http://www.nitrc.org/projects/conn).

The C++ core of the DMF code, together with Python and Octave/Matlab interfaces is publicly available at http://www.gitlab.com/concog/fastdmf.

The propofol and DOC patient data that support the findings of this study are available from Dr. Emmanuel Stamatakis, University of Cambridge upon reasonable request.

## Footnotes

1 Note that the excitatory and inhibitory populations within each region in the biophysical model are mutually and recursively coupled, and hence both excitation and inhibition are eventually affected by this procedure.

## Notes

### Competing Interest Statement

The authors have declared no competing interest.

### Summary of Updates

Updated link to FastDMF code repository.

